# Resolving ancient gene transfers clarifies the early co-evolution of eukaryotes and giant viruses

**DOI:** 10.1101/2023.07.11.548585

**Authors:** Sangita Karki, Frank O. Aylward

## Abstract

Members of the phylum *Nucleocytoviricota*, also called “giant viruses” due to their large physical dimensions and genome lengths, are a diverse group of dsDNA viruses that infect a wide range of eukaryotic hosts. Nucleocytoviruses likely evolved from smaller viruses, but the timing of their emergence and its relationship to the early evolution of eukaryotes remains unclear. Recent work has shown that the genomes of nucleocytoviruses often encode Eukaryotic Signature Proteins (ESPs) - including histones, vesicular trafficking factors, cytoskeletal components, and elements of RNA and DNA processing - that occur only rarely outside of eukaryotes. To investigate patterns of gene exchange between viruses and eukaryotes and possibly shed light on the early evolution of both, we examined the occurrence of viral-encoded ESPs (vESPs) and performed a comprehensive phylogenetic reconstruction on a subset that are widespread in nucleocytoviruses. Our results demonstrate that vESPs involved in cytoskeletal structure, ubiquitin system, and vesicular trafficking were acquired multiple times independently by nucleocytoviruses at different timepoints after the emergence of the eukaryotic supergroups. In contrast, vESPs involved in DNA and RNA processing are placed deep in their respective phylogenies, indicative of ancient gene exchange between nucleocytoviruses and eukaryotes. Examination of vESPs that could be rooted in archaea revealed that nucleocytoviruses likely acquired some of these genes prior to the emergence of the last eukaryotic common ancestor (LECA). Importantly, our findings also suggest that the eukaryotic delta DNA polymerase was acquired from nucleocytoviruses sometime during eukaryogenesis, underscoring the importance of viruses for early eukaryotic evolution. Collectively, these results suggest that gene exchange between nucleocytoviruses and eukaryotes played important roles in the evolution of both prior to the emergence of LECA.

## Introduction

The discovery of giant viruses, also known as nucleocytoplasmic large DNA viruses (NCLDVs), has dramatically expanded our understanding of the limits of viral size and genome complexity. These viruses belong to the phylum *Nucleocytoviricota* and comprise 6 orders, 11 families, and a rapidly-expanding number of family-level lineages [1]. Although they are found in diverse environments, they are particularly abundant in the oceans [2–5] and infect a wide range of hosts, ranging from the smallest known free-living eukaryote [6] to metazoans [7]. Members of families such as *Poxviridae*, *Asfarviridae*, and *Iridoviridae* generally include well-known vertebrate pathogens [8–10] while members of the *Mimiviridae, Phycodnaviridae,* and *Marseilleviridae* primarily infect protists and algae [11–13]. Phylogenomic studies across these groups have provided evidence that they have a shared evolutionary origin and potentially evolved from a tectivirus-like ancestor [14–16]. Gene acquisition from their hosts coupled with gene duplication likely contributed to the subsequent expansion of genome size and complexity in this lineage [14,17,18].

A resonant theme in recent years has been the discovery of numerous “cell-like” genes in nucleocytoviruses that are common in cellular life but rare or absent from other viral lineages. These include genes involved in translation [19,20], transcription [21], DNA replication and repair pathways [22–24], diverse metabolic pathways including the TCA cycle and glycolysis [25], cytoskeletal structure [26–28], membrane trafficking [29], and ubiquitin signaling [30]. In some cases viruses have acquired these genes from their hosts recently, and viral homologs form well-defined clades within broader eukaryotic lineages. This appears to be the case for ammonia transporters, some rhodopsins, and sphingolipid metabolism genes [31–33]. In other cases, such as histones and DNA polymerase subunits, viral genes are the result of ancient gene transfer events and their evolutionary relationship to cellular homologs is less clear [34,35]. Recent studies have shown that nucleocytoviruses often endogenize their genomes into those of their host [25,36,37] demonstrating that some of these genes may have originated in viruses and subsequently transferred to eukaryotes. Phylogenetic trees of multi-subunit RNA polymerase [21], DNA topoisomerase IIA [38], and actin [28] are examples of ancient virus-to-eukaryote gene transfers that have been proposed, although the direction of these transfer events is difficult to definitively ascertain. Regardless of the direction of gene transfer, these studies have begun to elucidate ancient patterns of gene transfer between viruses and eukaryotes that underscore the long co-evolutionary history of eukaryotes and nucleocytoviruses. This dynamic interplay between viruses and their hosts has shaped the early evolution of both and likely began prior to the emergence of the Last Eukaryotic Common Ancestor (LECA) [21].

The evolution of eukaryotes represents a major evolutionary transition in the evolution of life on Earth [39] and yet the details of this process remain a riddle [40]. Many complex features of eukaryotes emerged in the stem eukaryotic lineage prior to the emergence of LECA, but the order in which these traits emerged is largely unknown. Given the pre-LECA origin of nucleocytoviruses, it has been proposed that the co-evolution of these viruses and their hosts played a role in eukaryogenesis [41]. Interestingly, many nucleocytoviruses encode Eukaryotic Signature Proteins (ESPs) that are involved in central processes in eukaryotic cells, such as DNA and RNA processing, vesicular trafficking, cytoskeletal structure, and ubiquitin signaling. ESPs are common in eukaryotes but typically rare or absent in bacteria and archaea, and examination of their evolution therefore can provide key clues into the emergence of eukaryotes [42,43]. To provide deeper insight into the early co-evolution of nucleocytoviruses and eukaryotes we performed an in-depth survey of viral-encoded ESPs (vESPs) and performed a comprehensive phylogenetic reconstruction of their evolutionary histories. Our results provide key insights into the origin of nucleocytoviruses and the extent of virus-eukaryote gene transfer that occurred prior to the emergence of LECA.

## Results

### Occurrence of vESPs

We identified a set of 763 Pfam domains that are present in greater than 95 percent of the eukaryotic genomes surveyed, and 500 of these could be identified in at least one nucleocytovirus genome. These Pfams could be clustered into six groups depending on their occurrence across bacteria and archaea (S1 Data). Clusters 1 and 2 are broadly distributed across all cellular lineages (pan-cellular group), clusters 3 and 4 are found primarily in eukaryotes (true vESPs), and clusters 5 and 6 are found primarily in eukaryotes and archaea (archaea-vESPs) (Figure 1A). Many vESPs were only found in a small number of viral genomes and are not suitable for phylogenetic reconstruction; we therefore examined the prevalence of vESPs from different clusters across the *Nucleocytoviricota* to identify those that are broadly represented in this viral phylum (Figure 1B, 1C). To identify ESP that would be most likely to shed light on the early evolution of both nucleocytoviruses and eukaryotes, we searched both Pfams that were represented in greater than five nucleocytovirus genomes and were true ESPs (i.e., they belonged to clusters 3 or 4). A preponderance of these proteins are involved in vesicular trafficking, ubiquitin signaling, cytoskeletal structure, mRNA biogenesis, and DNA processing, and we therefore focused our analysis on protein families belonging to these functional categories. We focused our analysis on true vESPs, but in some cases we also included archaeal vESPs because of the opportunity to root these trees in the archaea.

**Fig 1.**
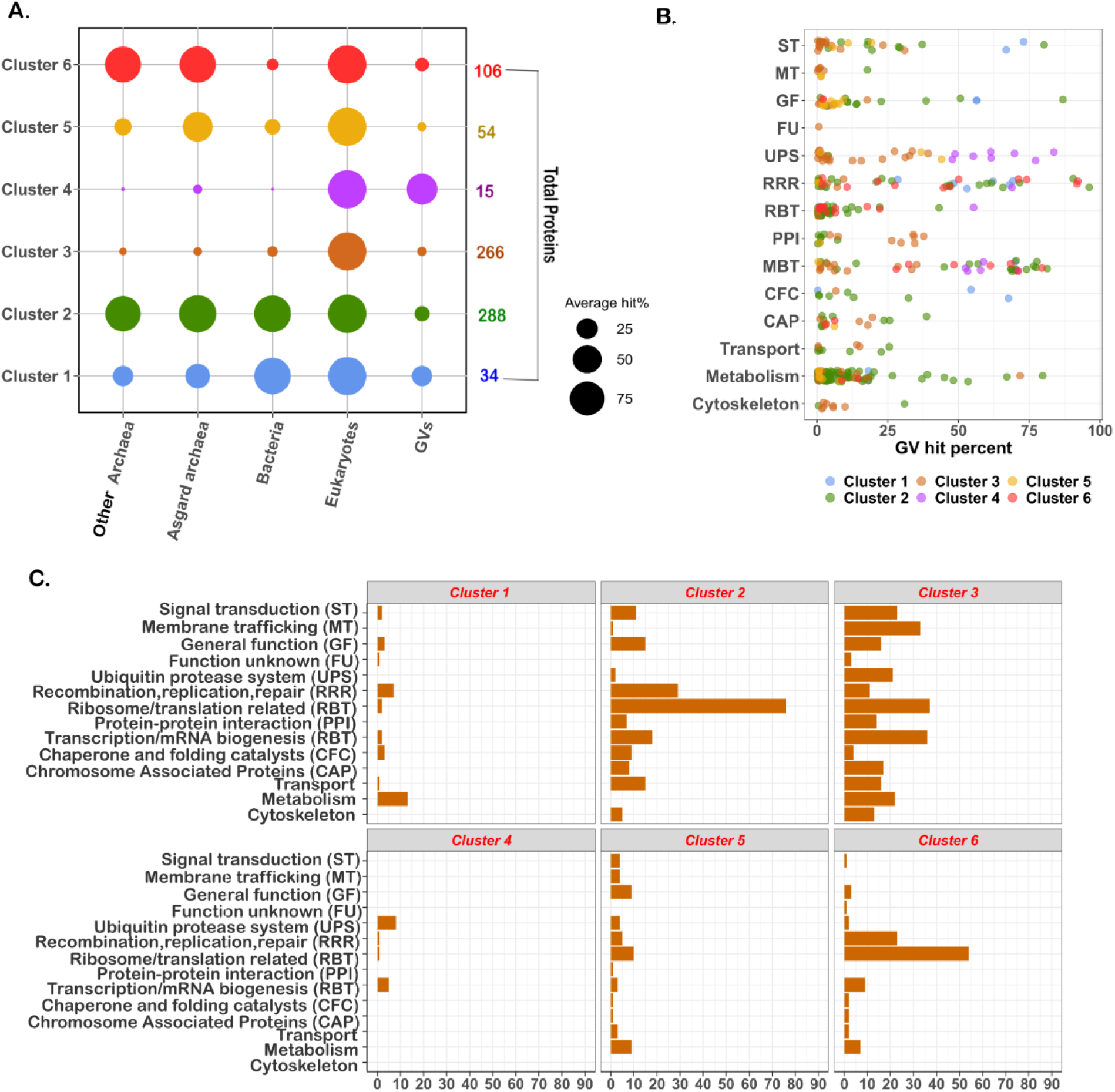
Clustering and functional annotation of ESPs. A) Unsupervised clustering of 763 eukaryotic proteins revealed six different clusters based on their prevalence in different groups. B) Functional categories of protein domains present in greater than 5 nucleocytovirus genomes irrespective of clusters. C) Functional annotation of protein domains belonging to different clusters.

Given the divergent nature of many vESPs that we examined here, we used several quality-control filters to examine the reliability of the trees that we generated. In order to evaluate the phylogenetic strength of the vESP trees we constructed, we calculated the Tree Certainty (TC) matric for each tree. This metric varies between zero and one, with higher values having fewer contrasting bipartitions and a more reliable topology [44]. Longer genes with a stronger phylogenetic signal typically exhibit larger TC values [45], and here we report only trees with TC > 0.5. All TC scores are provided for each tree and are also provided in S2 Data. In addition, for all vESPs we constructed trees using both the best fit site-homogeneous substitution model and the best fit complex model (i.e. site heterogeneous). The topologies of both trees were examined, and results are only reported when they are largely consistent (see Methods for details and S2 and S3 Text).

### Evolution of vESPs involved in ubiquitin signaling, cytoskeletal dynamics, and vesicular trafficking

Phylogenetic analysis of vESPs involved in ubiquitin signaling, vesicular trafficking, and cytoskeletal structure revealed topologies in which vESPs were nested within broader clades of eukaryotic proteins (Fig 2), clearly indicative of eukaryote-to-virus gene transfer events. It is not possible to root these trees given the lack of homologs in other cellular domains, but the placement of almost all vESPs within multiple distinct lineages nested within eukaryotic clades is strong evidence that viruses acquired these vESPs after the establishment of the main eukaryotic supergroups. Moreover, the occurrence of multiple distinct vESP clades demonstrates that viruses acquired most of these proteins multiple times independently throughout their evolution. It is also notable that the breadth of the vESP clades vary between trees. For example, whereas the SNARE and synaptobrevin vESPs are scattered broadly among eukaryotic homologs (Fig 2A), the Ub-activating enzyme and Ub-protease form some large clades (Fig 2B). Collectively this suggests that the vESPs were acquired by viruses over a range of different timescales, with shallow vESP clades representing recent acquisitions and deeper clades representing older gene transfer events.

**Fig 2.**
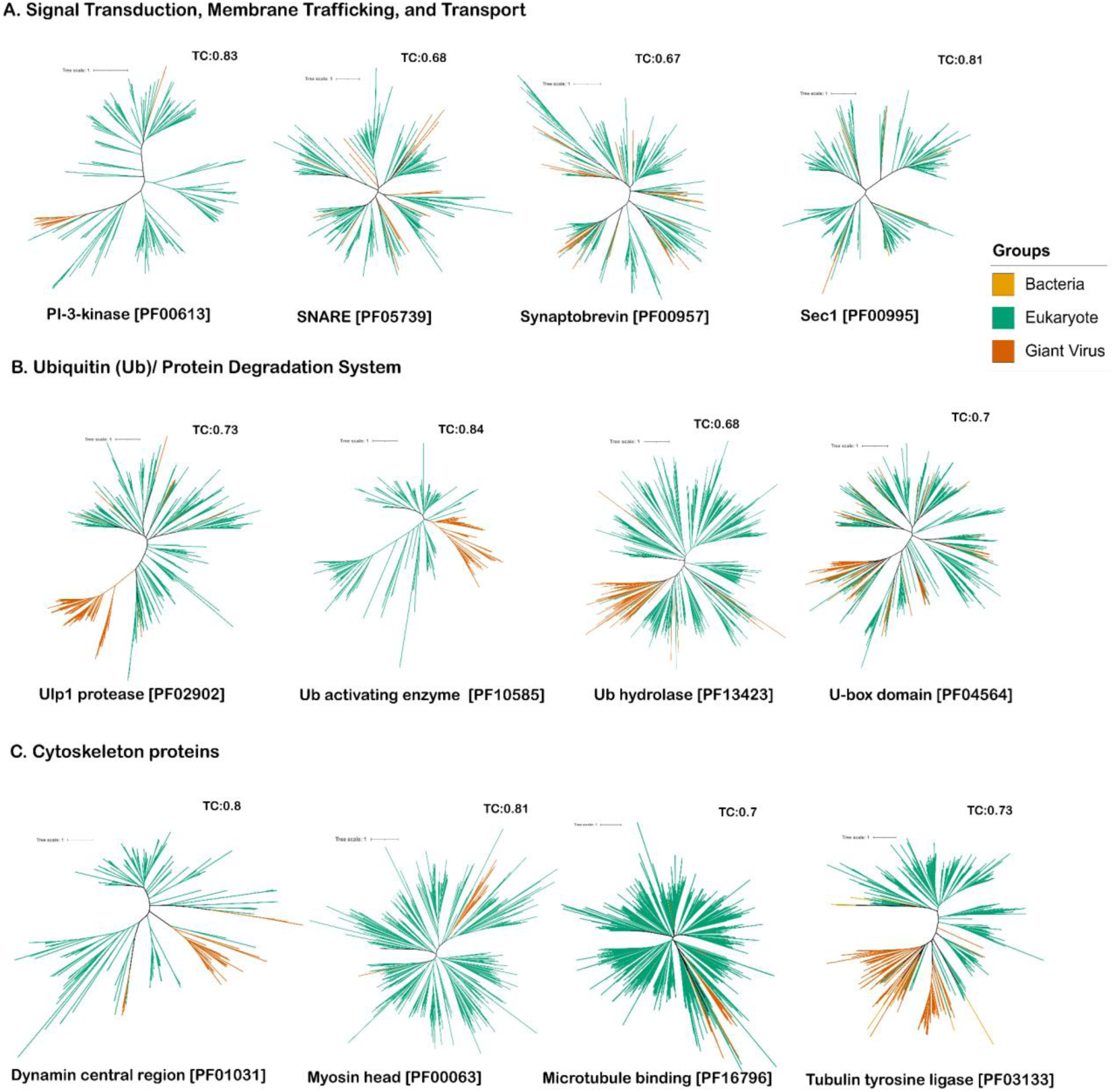
Phylogenetic trees for vESPs involved in (A) membrane trafficking, (B) ubiquitin system, and (C) cytoskeletal structure.

Our findings for a relatively recent viral acquisition membrane trafficking proteins is consistent with a phylogenetic study where a highly divergent Mimivirus Rab GTPase, a crucial regulator of membrane trafficking in eukaryotes, was reported to be acquired from unicellular eukaryotes [46]. Moreover, a recent study that confirmed the activity of some viral vesicular trafficking components also suggested that these proteins had been acquired multiple times by viruses [29]. Several phylogenetic studies of viral cytoskeleton homologs also suggested that the viral myosin and kinesin homologs form distinct clades nested within broader eukaryotic lineages [26,27]. Interestingly, a recent study of viral actin (i.e. viractin) homologs found evidence that viruses encoded this protein prior to its origin in modern eukaryotes [28], which stands in contrast to the general patterns we observed for cytoskeletal proteins. Viractin is relatively rare in nucleocytoviruses genomes and was therefore excluded from our analysis, but this previous study stands as a reminder that the relatively recent viral acquisition of eukaryotic cytoskeletal homologs may not hold for all protein families. Indeed, exceptions for these late branching clades were notable for some ubiquitin related proteins and a protein involved in membrane trafficking and biogenesis (ATG8) (Fig 1 in S1 Text).

### Phylogeny on DNA and RNA processing proteins belonging to true vESPs

The vESPs involved in mRNA and DNA processing formed deep-branching lineages in their respective trees, contrasting sharply with vESPs involved in other processes (Fig 3). This suggests that these proteins have a much more ancient evolutionary origin in the *Nucleocytoviricota*. Once again, it is not possible to root these trees due to the lack of homologs in other cellular domains, and it is therefore not possible to confidently determine the precise timing and direction of transfer that led to these topologies. Nonetheless, the unrooted trees still exhibit a stark distinction to those of vESPs involved in vesicular trafficking, cytoskeletal structure, and ubiquitin signaling, indicating that vESPs involved in mRNA and DNA processing have an ancient origin in the *Nucleocytoviricota* that can potentially be used to date the origin of this phylum.

**Fig 3.**
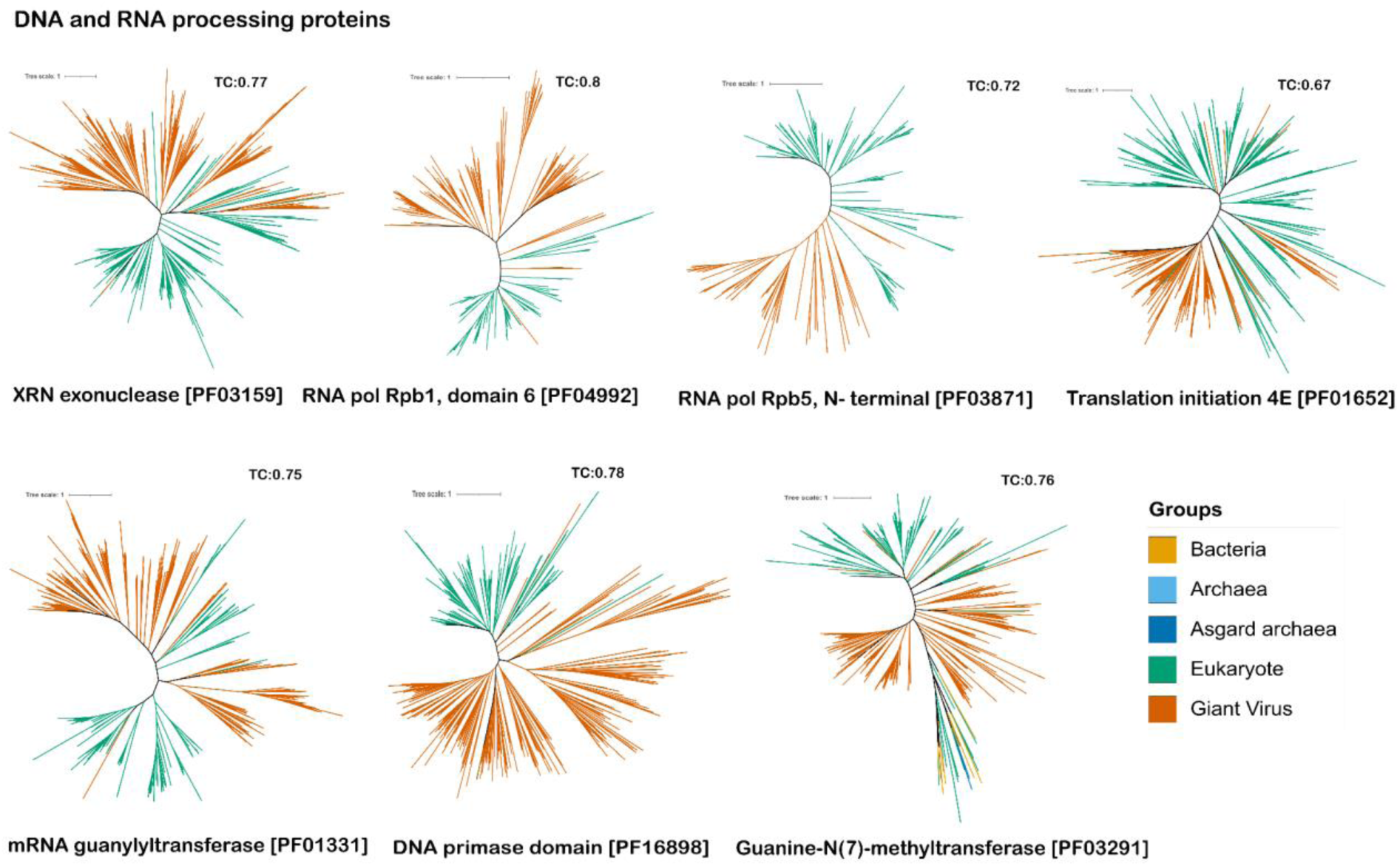
Phylogenetic trees for true vESPs involved in DNA and RNA processing.

Specifically, DNA primase, XRN endonuclease, different RNA polymerase subunits, enzymes of mRNA capping complex (mRNA cap guanylyltransferase, mRNA cap methyltransferase), and eukaryotic initiation factor 4E, all showed deep branching topology suggestive an ancient gene transfer between nucleocytoviruses and eukaryotes. Indeed, proteins belonging to the mRNA capping complex showed deep branching placement suggestive of ancient gene transfers between nucleocytoviruses and its hosts (Fig 3). Capping and methylation mechanisms are crucial steps in mRNA processing and initiation of protein synthesis for eukaryotes as well as giant viruses. A previous phylogenetic study on mRNA capping enzymes of nucleocytoviruses suggested that these two mRNA capping domains were present in the ancestor of nucleocytoviruses [47].

Translation initiation factor 4E (eIF4e) is involved in directing ribosomes to bind the cap structure of mRNA and therefore acts as a bridge between RNA processing and translation. Compared to other proteins involved in DNA or RNA processing, the tree of eIF4e provided strong evidence for multiple transfers of this protein from eukaryotes to viruses (Fig 3). Similar findings were reported in a previous study that broadly examined translation-associated genes in giant viruses and found that viral proteins were typically acquired multiple times independently from different eukaryotic lineages [18].

Overall, phylogenetic analysis of DNA and RNA processing protein domains belonging to cluster 3 and 4 (true vESPs) shows that these vESPs are mostly ancient. Although multiple viral clades were sometimes present in these trees, they were often placed deep in the tree in ancient nodes that are difficult to resolve. We therefore consider it more likely that this is a product of difficulties resolving deep nodes in a phylogeny rather than multiple gene transfer events. It is likely that the acquisition of core machinery for DNA and mRNA synthesis were critical for the early establishment of the *Nucleocytoviricota*, and it is therefore not surprising that these are the most ancient protein families in this phylum.

### Rooting gene transfer events with vESPs found in Archaea (Archaea-vESPs)

In order to infer direction of transfer, we expanded the analysis to include genes from the Archaea-vESP protein families because these can be rooted with the Archaea as an outgroup (Clusters 5 and 6 in Fig 1A). This is critical because it allows for a more precise evaluation of when giant viruses acquired these genes relative to the Last Eukaryotic Common Ancestor (LECA). Moreover, in some cases it has been proposed that early eukaryotes acquired certain genes from viruses prior to their diversification [21,28,38], and basal-branching placement of vESPs would provide evidence that this is a possibility. Although Archaea-vESPs are not true ESPs due to their presence in Archaea, the phylogenetic proximity of Asgard Archaea to early eukaryotes leaves open the possibility of gene transfer between some archaeal lineages and giant viruses during the early emergence of the *Nucleocytoviricota* [48]. Moreover, the Archaea-vESPs were enriched in genes involved in DNA and RNA processing, which provides the opportunity to examine how giant viruses acquired genes involved in these processes.

Similar to the phylogenies with the other vESPs, Archaea-vESPs involved in DNA and RNA processing exhibited deep branching viral clades indicative of ancient gene transfers with eukaryotes. Among the nine Archaea-vESPs we analyzed in detail, our rooted trees revealed basal-branching eukaryotic lineages in seven and basal-branching nucleocytovirus clades in two (Fig 4). The phylogeny of the RNA polymerase subunit 5-C terminal, which is common in Archaea and eukaryotes but absent in bacteria, revealed an early-branching viral clade. This suggests that this protein family already existed in the *Nucleocytoviricota* prior to the emergence of LECA, consistent with a previous study that focused on the two major RNA polymerase subunits and found that both of these protein families were found in the *Nucleocytoviricota* prior to the diversification of modern eukaryotes [21]. Similarly, a phylogeny of the DNA sliding clamp also revealed a deep branching viral clade indicative of an ancient presence in the *Nucleocytoviricota,* although several deep nodes in this tree had low bootstrap support (<80, Fig 9 in S2 and S3 Text) and must be interpreted with caution. For the other seven Archaea-vESPs, we recovered clear basal-branching eukaryotic clades with strong bootstrap support (S2 and S3 Text).

**Fig 4.**
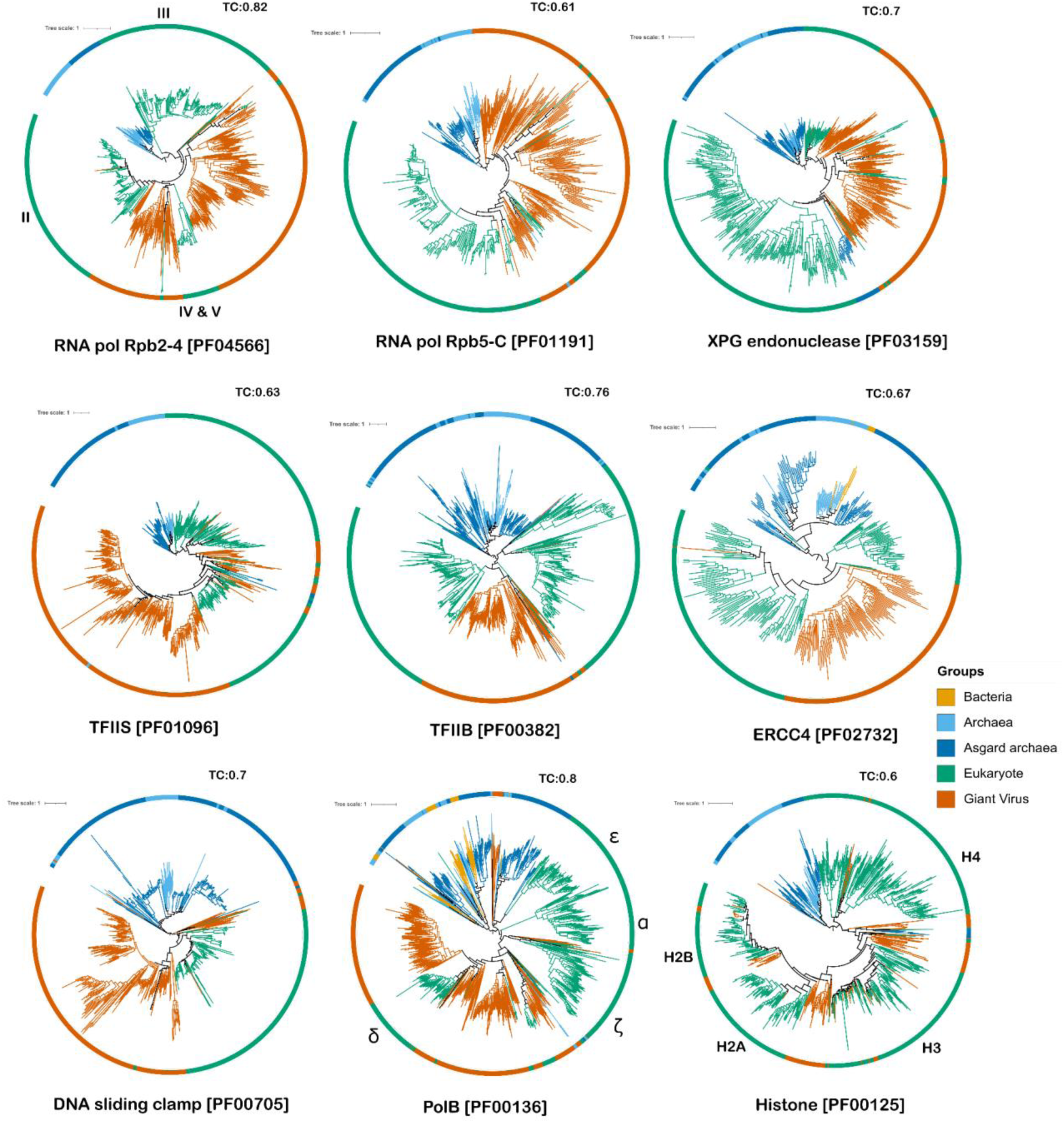
Phylogenetic trees for Archaea-vESPs involved in DNA and RNA processing.

Some protein families, such as multimeric RNA polymerase subunits and family B polymerases, diversified in eukaryotes through a complex process of duplication and divergence that took place prior to the emergence of LECA. It therefore remains a possibility that giant viruses acquired these proteins and then transferred them back to eukaryotes (i.e. reciprocal gene transfer), thereby participating in early diversification of these protein families. This has already been suggested for the two largest subunits of multimeric RNA polymerase, which are found in three copies in all modern eukaryotes but only a single copy in bacteria, archaea, and giant viruses. A previous study found that eukaryotic RNAP subunits form three distinct clades (I, II, and III), consistent with their early diversification in eukaryotes, and that RNAP II is nested within a clade of viral homologs. Our topology for RNAP subunit 2, domain 4, interestingly consists of II, III, IV&V clades and nonetheless suggests that giant viruses acquired RNAP before all three subunits had been established in eukaryotes (Fig 4). Similarly, we identified a case of reciprocal gene transfer for Transcription elongation factor IIS (C-terminal) which is responsible for regulating RNAP II, indicating that modern eukaryotes may have derived this key element of the transcriptional apparatus from nucleocytoviruses. The TC support values surrounding the nested eukaryotic clade are low (<0.6) and therefore should be interpreted with some caution, however.

Importantly, our phylogenetic analysis of family B DNA polymerases also suggests that reciprocal eukaryote-virus gene transfers have played a role in their evolution. Eukaryotes typically encode four main family B polymerases that were likely present in LECA [49]. These four PolBs form distinct clades in our tree, consistent with their ancient origin. Interestingly, the eukaryotic delta family PolB (PolDelta) clade - which includes representatives from all major eukaryotic supergroups-is clearly nested within the major PolB clade of the *Nucleocytoviricota*. This strongly suggests that the modern eukaryotic PolDelta subunit is derived from an ancient acquisition from giant viruses that took place during the early diversification of the PolB family in eukaryotes. Several other smaller eukaryotic clades are nested within the *Nucleocytoviricota* in this tree, demonstrating that more recent transfers from viruses to eukaryotes have further complicated the evolutionary history of this protein family. This is consistent with the hypothesis presented in an early comparative genomic study that suggested eukaryotic PolDelta arose through interactions with viruses [35]. Similarly, the phylogeny of DNA topoisomerase IIA showed acquisition of this gene by the eukaryotic host from nucleocytoviruses (S1 Text, Fig 1), consistent with a previous study which showed viral origin of eukaryotic DNA topoisomerase IIA [38]. Together with the possible pre-LECA origin of the DNA sliding clamp, which associates with PolDelta, these results suggest that the DNA replication machinery of the Nucleocytoviricota originated prior to the diversification of the major eukaryotic supergroups.

One last example of possible eukaryote-virus reciprocal gene transfer involves the core histones. All modern eukaryotes have four major types of histones (H2A, H2B, H3 and H4), all of which have also been found in giant viruses [34]. Phylogenetic analysis of core histone proteins revealed the presence of viruses near the base of the major core histone diversification, consistent with a previous study that suggested that giant viruses had acquired these protein families prior to the emergence of LECA [50]. The overall topology of this tree suggests that multiple eukaryotes to virus gene transfers occurred, and it is possible that some of the eukaryotic clades arose through acquisition of viral homologs. The broad distribution of viral histone homologs, together with the overall low TC value for this tree (0.6) somewhat obscures these conclusions, however (Fig 4).

### Comparison with the *Mirusviricota*

A recent study discovered a novel phylum of large DNA viruses referred to as the *Mirusviricota* that has evolutionary links to the Nucleocytoviricota. Mirusviruses encode a herpesvirus-like morphogenetic module but otherwise have many components of DNA and RNA processing machinery that resemble nucleocytoviruses, and it remains unclear which lineage emerged first. We therefore conducted separate phylogenetic analysis of several vESPs that are also found in mirusviruses, including a DNA primase, RNA polymerase, TFIIS, ERCC4, PolB, DNA sliding clamp, and core histones. Our phylogenetic interpretation of these genes did not change when mirusviruses were included. Moreover, in most cases mirusvirus sequences clustered clearly within broader clades of nucleocytoviruses, with the exception of DNA topoisomerase IIA and DNA primase (Fig 2 in S1 Text).

### Quantitative comparison of tree depth across vESP categories

In order to quantitatively compare the timing at which different genes emerged in the *Nucleocytoviricota*, we examined ultrametric trees of all protein families analyzed in this study and identified the depth at which the first appeared in this viral phylum. The true vESPs trees were midpointed while Archea-vESPs were archeal rooted, and scenarios in which the most basal-branching lineage was viral were given a depth of zero (Fig 5). As expected, the DNA and RNA processing vESPs were the most ancient. Our tree depth analysis showed that a giant viral clade of mRNA cap guanylyltransferase, DNA primase, DNA sliding clamp as well as RNAP subunits are at the root, indicative of an ancient viral origin. By comparison, vESPs involved in cytoskeletal structure, vesicular trafficking, and ubiquitin signaling appeared in the *Nucleocytoviricota* more recently (Fig 5).

**Fig 5.**
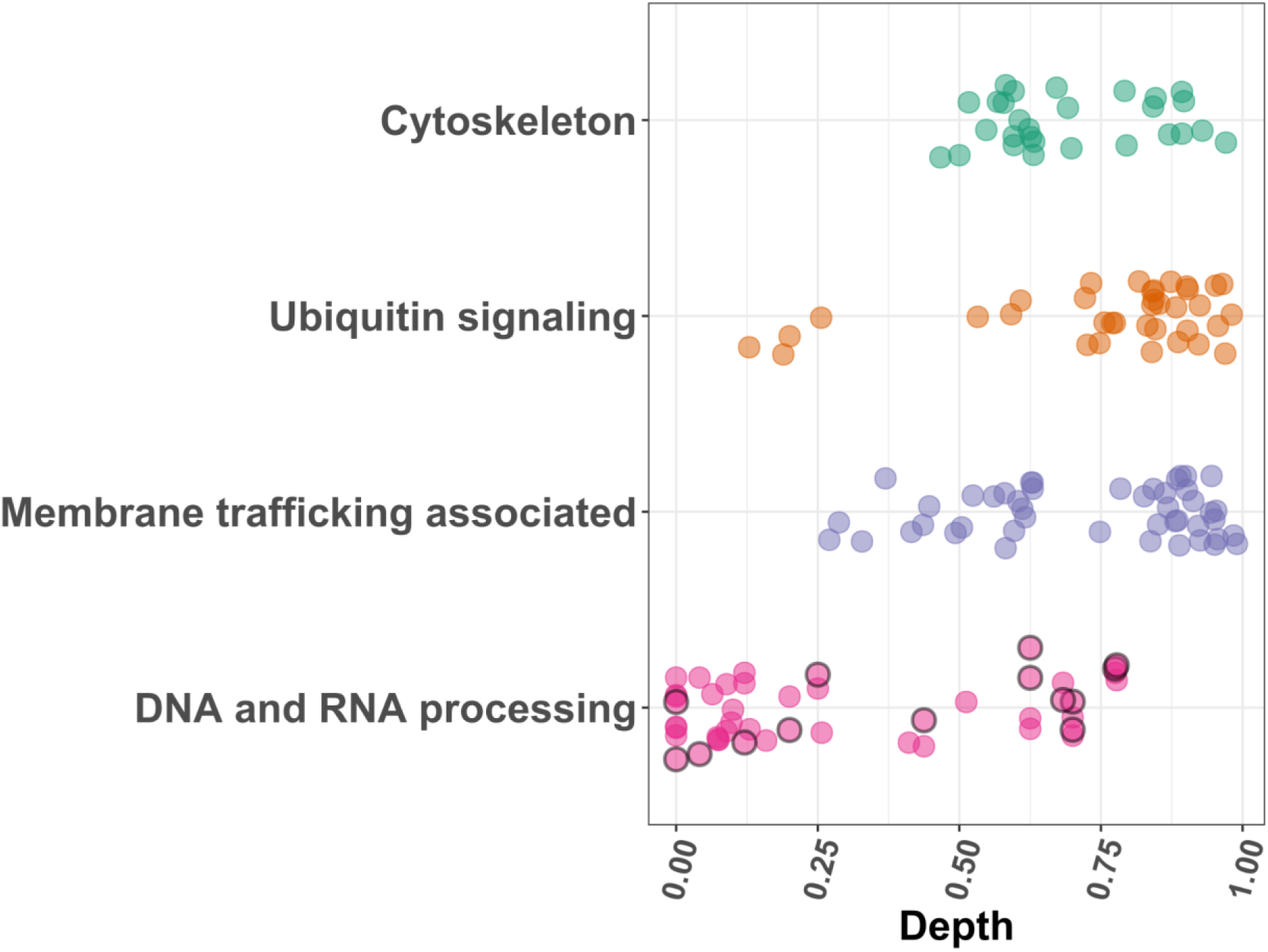
Phylogenetic tree depth for Membrane trafficking, Ubiquitin system, Cytoskeleton, and DNA and RNA associated proteins.

1. X-axis value 0 indicates root while 1 indicates tip in the phylogenetic tree and 2. Black border around point on DNA and RNA processing proteins represent Archaea-vESPs.

## Discussion

In this study we present a comprehensive phylogenetic analysis of ESPs encoded in nucleocytoviruses to evaluate virus-host gene transfers that may have occurred early in the evolution of eukaryotes. Phylogenies on cytoskeletal proteins, membrane trafficking associated proteins, and ubiquitin signaling proteins generally showed late branching topologies indicative of multiple recent viral acquisitions. In contrast, phylogenies of DNA and RNA processing vESPs revealed deeper branching topologies indicative of more ancient gene exchange between the eukaryotes and giant viruses.

A defining feature of the *Nucleocytoviricota* is the virus factory, also called the viroplasm, which is a membrane-bound intracellular structure that forms during infection and is the location of DNA synthesis, viral transcription, and virion morphogenesis. Although not all members of the *Nucleocytoviricota* form virus factories during infection, it is a prevalent feature of giant viruses spanning all six orders and two classes of this phylum and is most likely a trait that was present in their last common ancestor. Central importance has been placed on virus factories in the biology of nucleocytoviruses [51], and any study that seeks to clarify the origin of the *Nucleocytoviricota* must therefore consider the origin of this viral structure. Proper functioning of the virus factory would necessarily require a viral mechanism for DNA polymerization, transcription, and mRNA processing, and it is therefore fascinating to note that vESPs involved in these processes exhibit the most ancient evolutionary origins of any proteins in the *Nucleocytoviricota*. We therefore propose that the most parsimonious scenario that explains the deep-branching topologies the family B DNA polymerase, DNA sliding clamp, topoisomerase, multimeric RNA polymerase, and mRNA capping machinery is pre-LECA origin of the virus factory. Recent evidence suggests that stem Eukaryotes diversified for a long period of time prior to the emergence of LECA [52], and it is therefore not only plausible but likely that these stem Eukaryotes housed a complex array of viruses that gave rise to the modern *Nucleocytoviricota* [53].

It is possible that the virus factory was a key innovation that potentially allowed early nucleocytoviruses to evade host defenses by creating a physical barrier between the host cytoplasm and the site of viral replication and transcription. This is analogous to the infection strategy used by many large bacteriophages [54], and is therefore a mechanism employed by several distinct lineages in the virosphere. It is unclear whether early eukaryotes possessed a fully developed nucleus at the time that the virus factory originated, and it is possible that the emergence of the virus factory pre-dates the nucleus. Due to the mechanistic similarity of nuclei and virus factories, it has been proposed the former may have evolved from the latter [55,56]. While our analysis cannot confirm or refute this hypothesis, evidence for a viral origin of eukaryotic machinery involved in DNA replication and transcription suggests that nucleocytoviruses played a role in the emergence of modern eukaryotes.

A recent study discovered a novel phylum of large DNA viruses, the *Mirusviricota*, that are related to the *Nucleocytoviricota* but encode a herpesvirus like capsid [57]. The Mirusviricota encompass a broad phylogenetic diversity and it has been suggested that they may represent a progenitor to the *Nucleocytoviricota*. According to this scenario, nucleocytoviruses potentially emerged through loss of the herpesvirus-like capsid and acquisition of a double-jellyroll major capsid protein and other associated structural components. To examine this possibility, we included Mirusviruses in the trees of DNA and RNA processing vESPs, which are the most ancient protein families in the Nucleocytoviricota (Fig 2 in S1 Text). In most cases the *Mirusviricota* formed well-supported clades within the *Nucleocytoviricota*, providing evidence that the latter have a more ancient evolutionary origin and that the *Mirusviricota* likely emerged from within the *Nucleocytoviricota*. We still know little of the enigmatic origins of the *Mirusvirocita* due to their recent discovery and a paucity of studies that have examined this lineage, however the evolutionary origins of the *Mirusviricota* should therefore be revisited as more genomes become available.

Although many vESPs involved in DNA and RNA processing have likely pre-LECA origins in the *Nucleocytoviricota*, this is not the case for all protein families in these categories. For example, the topologies of XPG and ERCC4 endonucleases genes clearly have basal-branching eukaryotic lineages that suggests nucleocytoviruses acquired these genes after the emergence of the major eukaryotic supergroups. We therefore propose that the viral acquisition of cellular genes was a stepwise process that took place in multiple stages (Fig 6). vESPs critical for virus factory functioning were acquired first at the pre-LECA origin of the *Nucleocytoviricota* (stage I), and further acquisition of additional genes involved in DNA and RNA processing and transcriptional regulation continued shortly after the emergence of modern eukaryotic diversity (stage II). In the third stage, multiple distinct lineages of nucleocytoviruses independently acquired genes involved in cytoskeletal dynamics, ubiquitin signaling, vesicular transport, and a wide range of other processes. Collectively these stages explain both the evolutionary origins of the *Nucleocytoviricota* as well as the subsequent emergence of remarkably large and complex genomes within multiple lineages of this group. Moreover, these findings highlight examples of reciprocal gene exchange between viruses and early eukaryotes and provide strong evidence for a viral origin of the eukaryotic family B DNA polymerase PolDelta. Our results underscore the importance of understanding cellular evolution in the context of co-evolution between cells and viruses.

**Fig 6.**
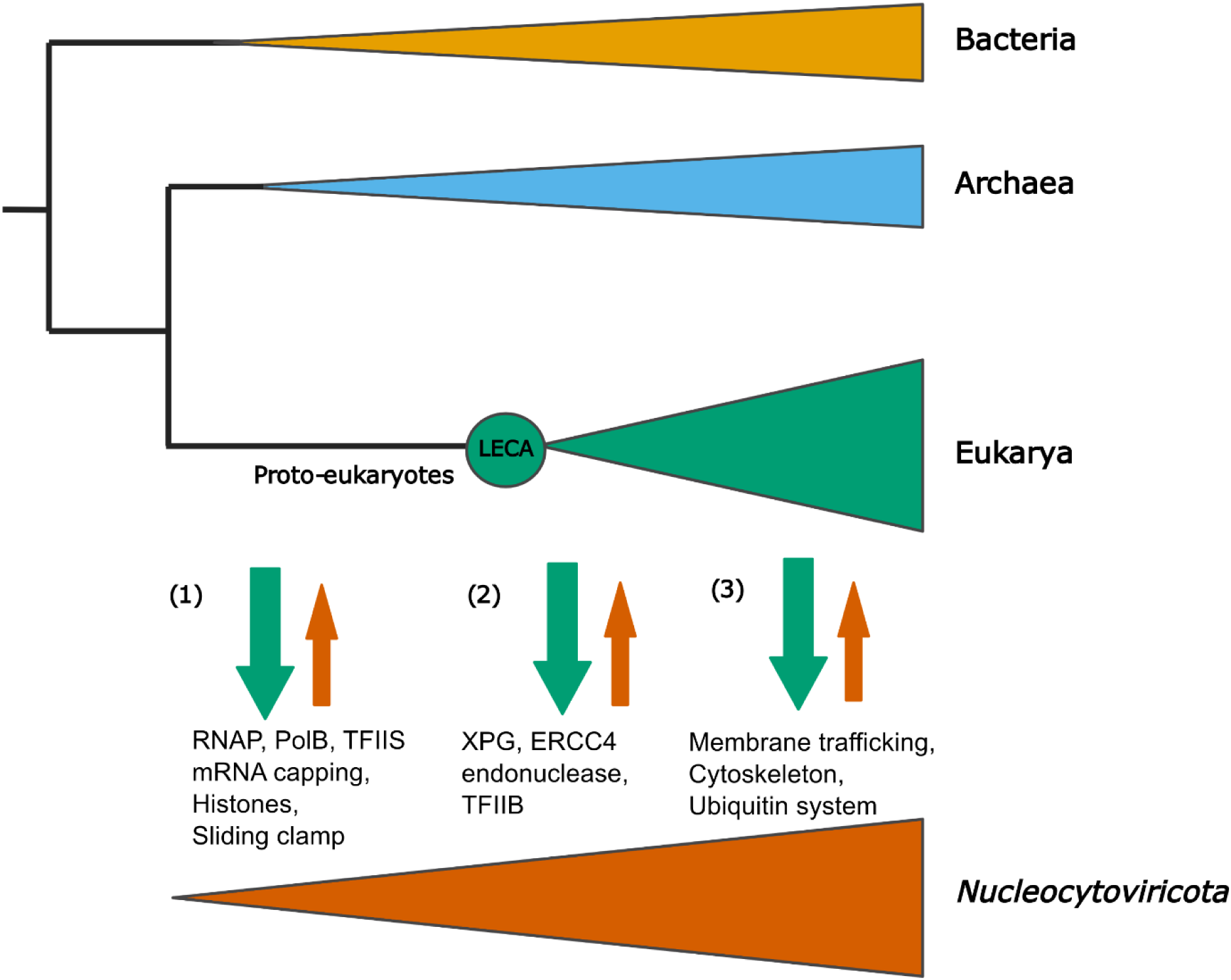
Schematic diagram showing gene exchange between the eukaryotes and giant viruses. Our results indicate three scenarios (1) gene exchange between ancient giant viruses and proto-eukaryotes (2) gene exchange between giant viruses and eukaryotes post-LECA (3) recent gene exchange.

## Materials and Methods

### Compilation of genomic datasets used in this study

Prior to phylogenetic analysis we compiled a set of eukaryotic, archaeal, bacterial, and nucleocytovirus genomes. We downloaded high-quality eukaryotic genomes from the eggNOG v5.0 database. Because eggNOG has relatively few protist genomes, we also included complete and chromosome-level genomes for select protists available on the National Center for Biotechnology Information (NCBI) databases as of October 8, 2021. For bacterial and archaeal genomes, we retrieved genomes from the Genome Taxonomy Database (GTDB, v95). To enrich our database in asgard archaeal genomes, we also included Asgard archaea genomes as reported in [58] that were not already present in the GTDB. For giant viruses, we used a set of curated genomes available on the Giant Virus database (https://faylward.github.io/GVDB/) that have been described previously [1]. For mirusviruses, we used genomes that were recently reported [57]. For eukaryotic genomes we used protein predictions available on EggNOG, and for all other taxa we predicted proteins using Prodigal v. 2.6.3 with default parameters [59]. A full list of all genomes used is available on https://doi.org/10.5281/zenodo.8128475.

### Eukaryotic Signature Proteins (ESPs) detection, Clustering, and Functional annotation

For *de novo* identification of ESPs, we searched bacterial, archaeal, viral, and eukaryotic genomes against the Pfam database and searched for domains that were enriched in eukaryotes. For this we used the hmmsearch command in the HMMER3 v.3.3 package with the “--cut_nc” parameter [60], with the Pfam database version 32.0 [61] as a query. To determine which Pfams were ESPs, we first removed all Pfams that were not present in >95% of all eukaryotic genomes, leaving us with a set of 763 protein families (See S1 Data). We then sought to cluster these Pfams into groups based on their distribution across the taxonomic groups in our analysis. For this we employed an unsupervised learning approach using Self-Organising Maps (SOMs) in R version 3.6.3, implemented with the kohonen package (script for this is available in https://github.com/sangitakarki/Self-Organising-Maps). To choose the optimal number of clusters we used the heuristic “elbow method”, and ultimately arrived at six groups (Fig 1A). To determine the role of these eukaryotic proteins we performed functional annotation of proteins using hmmsearch with 1e-5 threshold against the KEGG database (KEGG release 99.0, ver. 2021-08-01) [62]. Since the same genes had hits to multiple KEGG categories, we manually curated the results from the KEGG annotation to refine categories.

### Phylogenetic Analysis and Evaluation of Complex Models

For phylogenetic reconstruction, we retrieved protein sequences with hits to the Pfams under investigation, aligned them together using Clustal Omega v1.2.4 with default parameters [63], and used trimAl v1.4. rev15 for alignment trimming (parameter -gt 0.1) [64]. We then reconstructed maximum likelihood phylogenetic trees using IQtree v1.6.12 [65] with the option -bb 1000 to generate 1,000 ultrafast bootstraps [66], -m MFP to determine the best-fit model [67], -nt AUTO and --runs 5 to select the highest likelihood tree. Several best fit substitution models for different trees were selected based on Bayesian Information Criterion (BIC) [67] (full list of best fit models is reported in S3 Data). Most DNA and RNA processing protein trees were inferred under the LG [68] and VT [69] amino-acid exchange rate matrices, amino-acid frequencies (F), and variation in evolutionary rates across sites (R) [70], with different numbers of categories. Because amino acid substitution rates likely vary across alignments, we also inferred trees using complex models (C-models) that have different substitution matrices for every position in the alignment [71]. Since running complex models is time and memory consuming we used a new posterior mean site frequency (PMSF) model [72] to compare our models from -MFP option to C-models (C10 to C60). Although the trees inferred with the -MFP option generally had lower BICs, we still examined the trees inferred with complex models to assess any differences in topology that could be detected using the different methods. In almost all cases, we observed similar topologies between the two methods. One notable example is RNA pol subunit 5C, for which different groups of nucleocytoviruses were placed as basal branching when comparing the trees inferred with different methods (S3 Text).

All trees were visualized using the Interactive Tree of Life (iTOL) [73]. If we observed long branches that potentially represented rogue taxa, we removed these sequences from the analysis and re-performed alignment, trimming, and phylogenetic reconstruction. In some cases, the number of viral proteins outnumbered those of all other taxa by a factor of two or more. In these cases, we downsampled the viral proteins by choosing one representative nucleocytoviral genome from each family, with the genome with the highest N50 contig size chosen as the representative. This was done on the grounds that a higher N50 contig size is indicative of a higher quality genome assembly. In some cases, down sampling was also done for bacteria and archaea to class level and eukaryotes and Asgard archaea to respective representatives.

### Assessing Tree Quality and Tree Depth

In order to evaluate the phylogenetic congruency of the trees we used the Tree Certainty (TC) metric, which provides a measure of overall tree quality [44]. This was done in RaxML version 8.2.12 [74] (options -t and -z used to input the tree file and bootstrap file, respectively). All TC scores along with alignment length are available in S2 Data. We calculated the depth of viral clades in trees using a script that utilizes ape package [75] in R, bio-python and python.

## Data Availability

All genomes, proteins, and alignment used in this study can be found here: https://doi.org/10.5281/zenodo.8128475

All phylogenetic trees are available on iTOL using the username sangita1: https://itol.embl.de/shared_projects.cgi

All codes used in this study can be found here: https://github.com/sangitakarki/NCLDV_coevolution

## Acknowledgments

We acknowledge the use of the Virginia Tech Advanced Research Computing Center for bioinformatic analyses performed in this study. We are also thankful to the members of Aylward lab for their helpful suggestions.

## Supporting information

### S1 Text. Supporting figures

Fig 1. Phylogeny for proteins belonging to DNA processing, ubiquitin signaling, transport/trafficking and cytoskeleton dynamics respectively. Figure 2. Phylogeny with Mirusvirus for DNA and RNA processing proteins using a complex model (C60+F+G).

### S2 Text. Supporting figures

Fig 1: ERCC4 domain phylogeny reconstructed using VT+F+R8 model (generated by -MFP parameter) with bootstrap support value represented by black dots (bootstrap > 80% is shown).

Fig 2: Histone phylogeny reconstructed using VT+R8 model (generated by -MFP parameter) with bootstrap support value represented by black dots (bootstrap > 80% is shown).

Fig 3: Polymerase family B phylogeny reconstructed using VT+F+R10 model (generated by - MFP parameter) with bootstrap support value represented by black dots (bootstrap > 80% is shown).

Fig 4: XPG endonuclease phylogeny reconstructed using VT+F+R10 model (generated by -MFP parameter) with bootstrap support value represented by black dots (bootstrap > 80% is shown).

Fig 5: Transcription factor TFIIB phylogeny reconstructed using LG+R8 model (generated by - MFP parameter) with bootstrap support value represented by black dots (bootstrap > 80% is shown).

Fig 6: Transcription factor TFIIS phylogeny reconstructed using VT+F+R8 model (generated by -MFP parameter) with bootstrap support value represented by black dots (bootstrap > 80% is shown).

Fig 7: Phylogeny for RNA pol Rpb2, domain 4 reconstructed using LG+F+R10 model (generated by -MFP parameter) with bootstrap support value represented by black dots (bootstrap > 80% is shown).

Fig 8: Phylogeny for RNA pol Rpb5, C terminal reconstructed using VT+F+R7 model (generated by -MFP parameter) with bootstrap support value represented by black dots (bootstrap > 80% is shown).

Fig 9: Phylogeny for DNA sliding clamp reconstructed using LG+F+R8 model (generated by - MFP parameter) with bootstrap support value represented by black dots (bootstrap > 80% is shown).

### S3 Text. Supporting figures

Fig 1: ERCC4 domain phylogeny reconstructed using a complex model (C60 + F +G) with bootstrap support value represented by black dots (bootstrap > 80% is shown).

Fig 2: Core Histone phylogeny reconstructed using a complex model (C60 + F +G) with bootstrap support value represented by black dots (bootstrap > 80% is shown).

Fig 3: Polymerase family B phylogeny reconstructed using a complex model (C60 + F +G) with bootstrap support value represented by black dots (bootstrap > 80% is shown).

Fig 4: XPG endonuclease phylogeny reconstructed using a complex model (C60 + F +G) with bootstrap support value represented by black dots (bootstrap > 80% is shown).

Fig 5: Transcription factor TFIIB phylogeny reconstructed using a complex model (C60 + F +G) with bootstrap support value represented by black dots (bootstrap > 80% is shown).

Fig 6: Transcription factor TFIIS phylogeny reconstructed using a complex model (C60 + F +G) with bootstrap support value represented by black dots (bootstrap > 80% is shown).

Fig 7: Phylogeny for RNA pol Rpb2, domain 4 reconstructed using a complex model (C60 + F +G) with bootstrap support value represented by black dots (bootstrap > 80% is shown).

Fig 8: Phylogeny for RNA pol Rpb5, C-terminal reconstructed using a complex model (C60 + F +G) with bootstrap support value represented by black dots (bootstrap > 80% is shown).

Fig 9: Phylogeny for PCNA reconstructed using a complex model (C60 + F +G) with bootstrap support value represented by black dots (bootstrap > 80% is shown).

**S1 Data. Clusters, annotations, genome hits and other statistics for 763 proteins reported in this study.**

**S2 Data. TC scores and alignment length for phylogeny presented in this study.**

**S3 Data. Lists of site homogenous best fit models according to Bayesian criteria for phylogeny presented in this study.**

