## Supplementary material for "Resolving ancient gene transfers clarifies the early co-evolution of eukaryotes and giant viruses": S1 Text

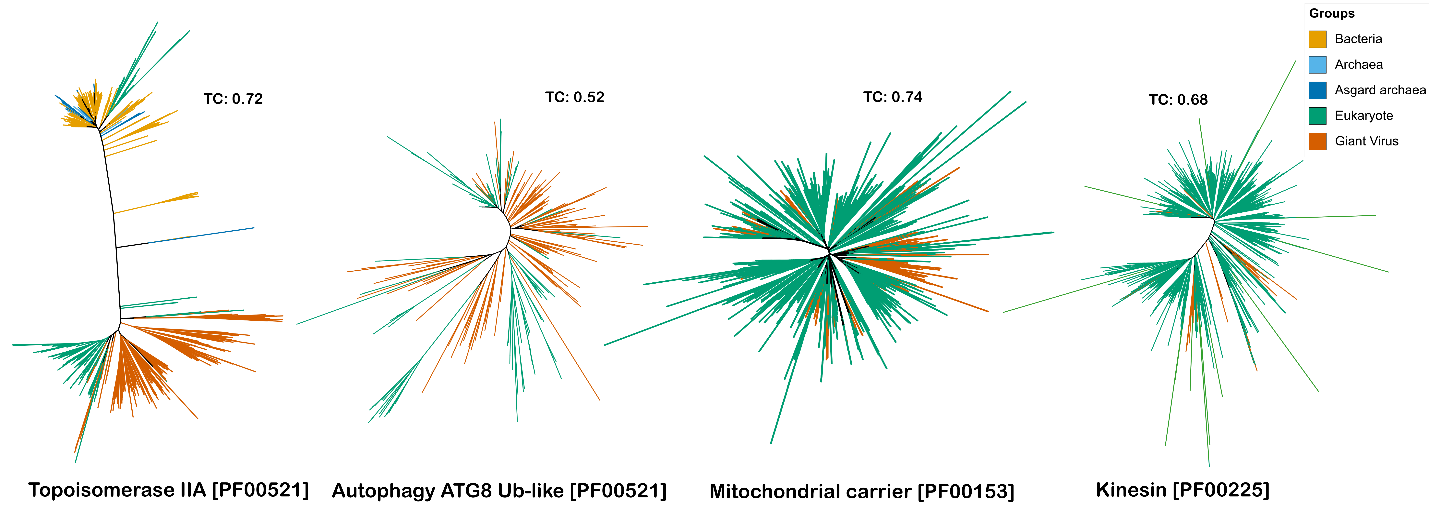

Fig 1. Phylogeny for proteins belonging to DNA processing, ubiquitin signaling, transport/trafficking and cytoskeleton dynamics respectively.

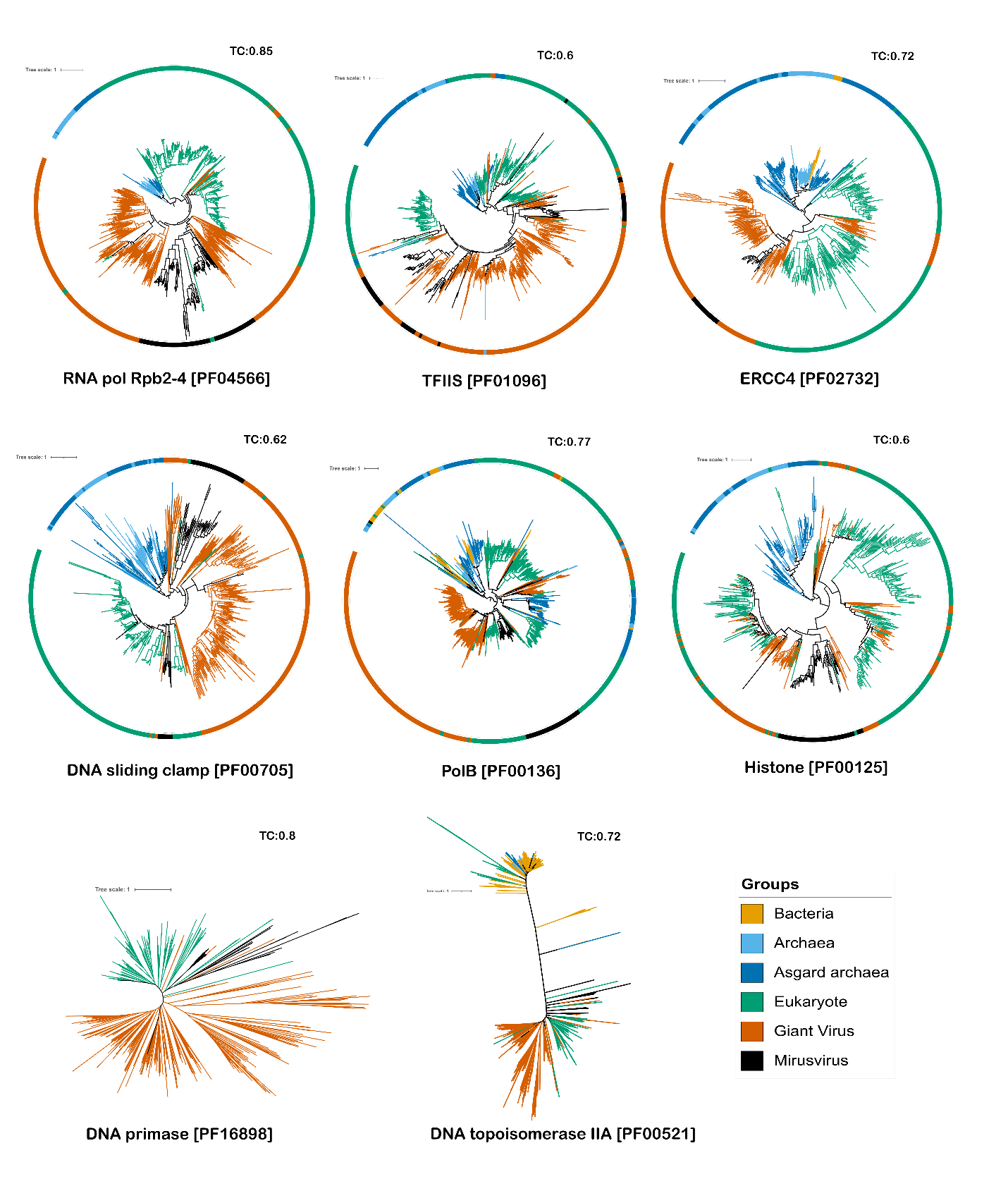

Figure 2. Phylogeny with Mirusvirus for DNA and RNA processing proteins using a complex model (C60+F+G).
