## Supplementary material for "Resolving ancient gene transfers clarifies the early co-evolution of eukaryotes and giant viruses": S2 Text

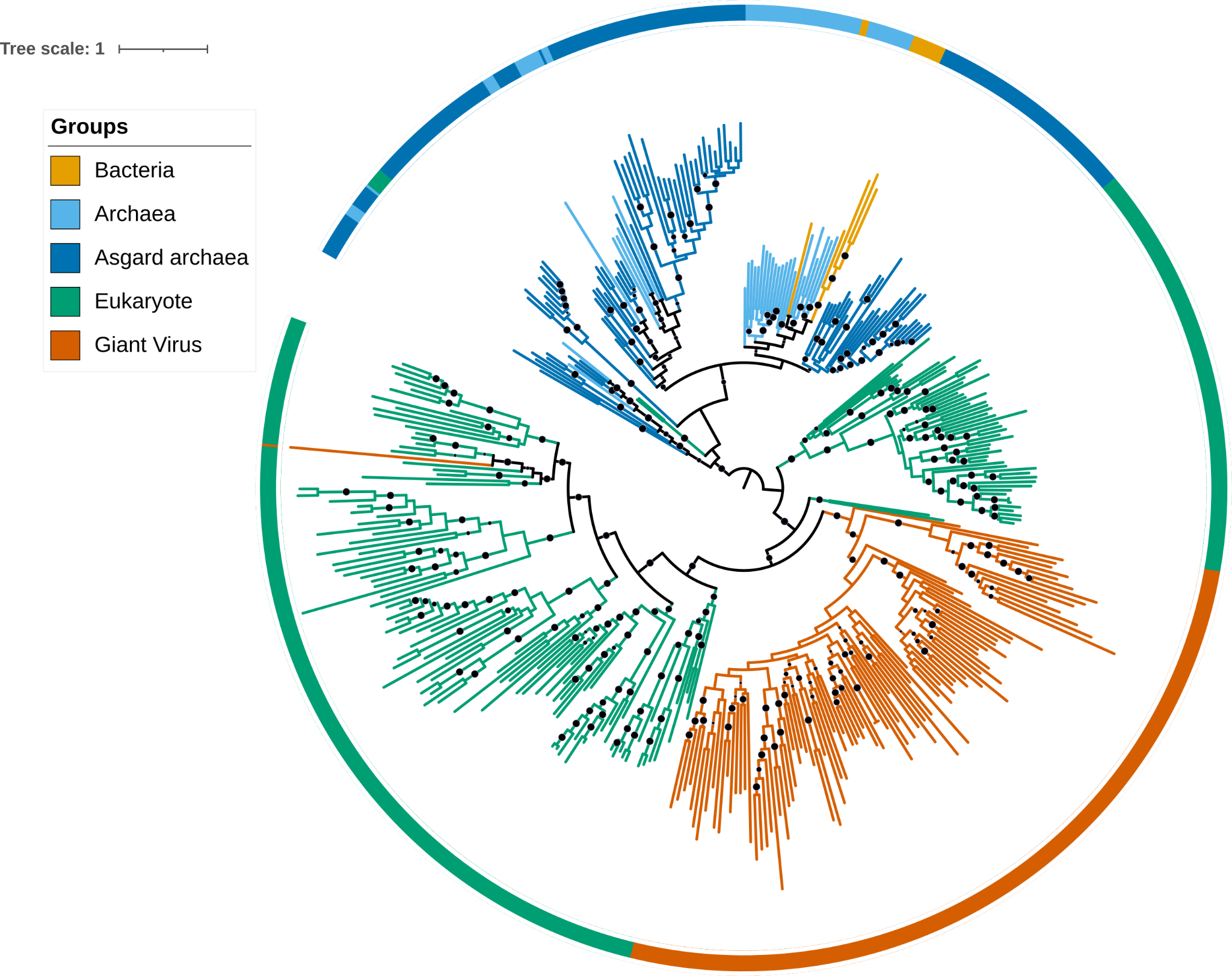

Figure 1: ERCC4 domain phylogeny reconstructed using VT+F+R8 model (generated by -MFP parameter) with bootstrap support value represented by black dots (bootstrap > 80% is shown).

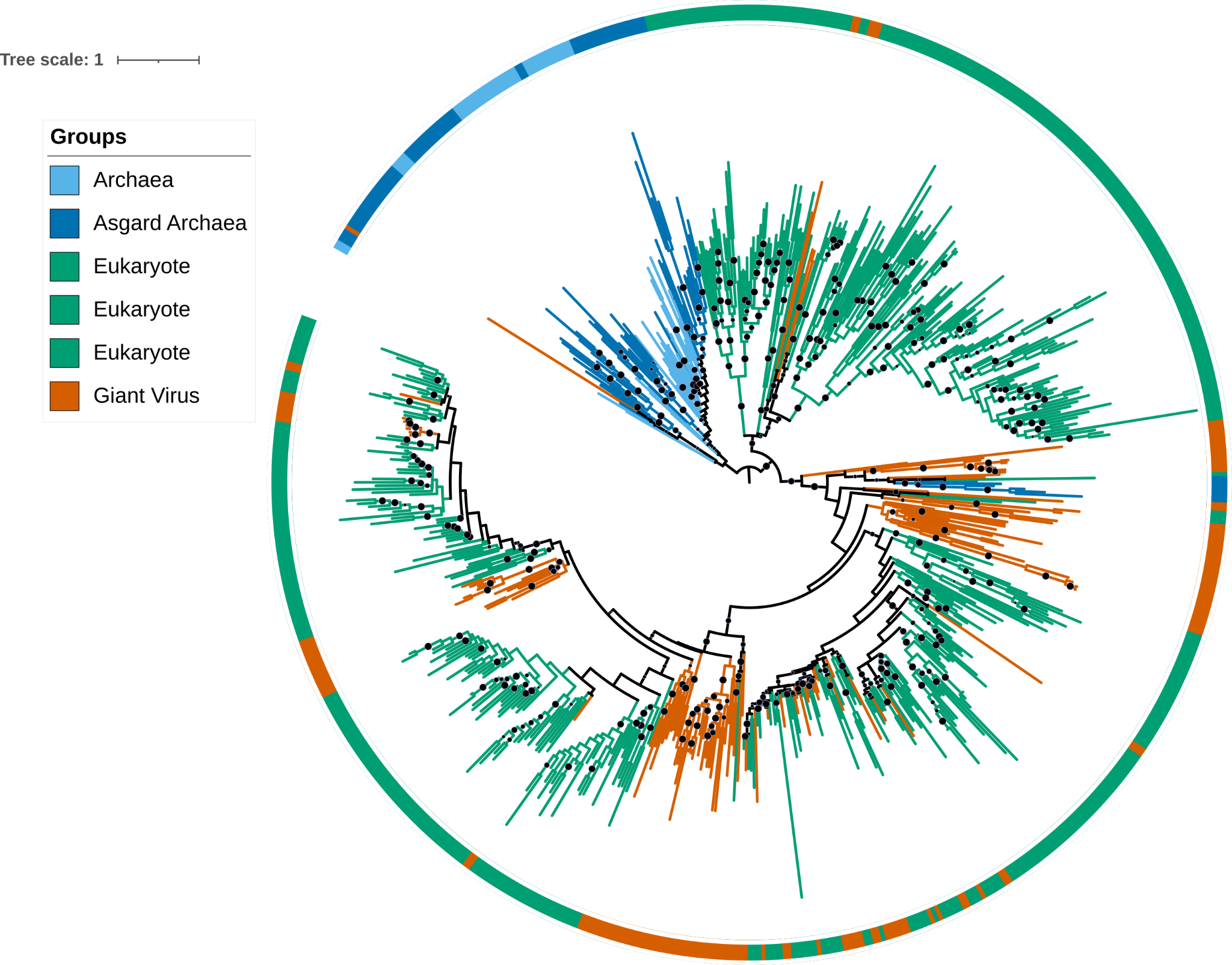

Figure 2: Histone phylogeny reconstructed using VT+R8 model (generated by -MFP parameter) with bootstrap support value represented by black dots ( bootstrap > 80% is shown).

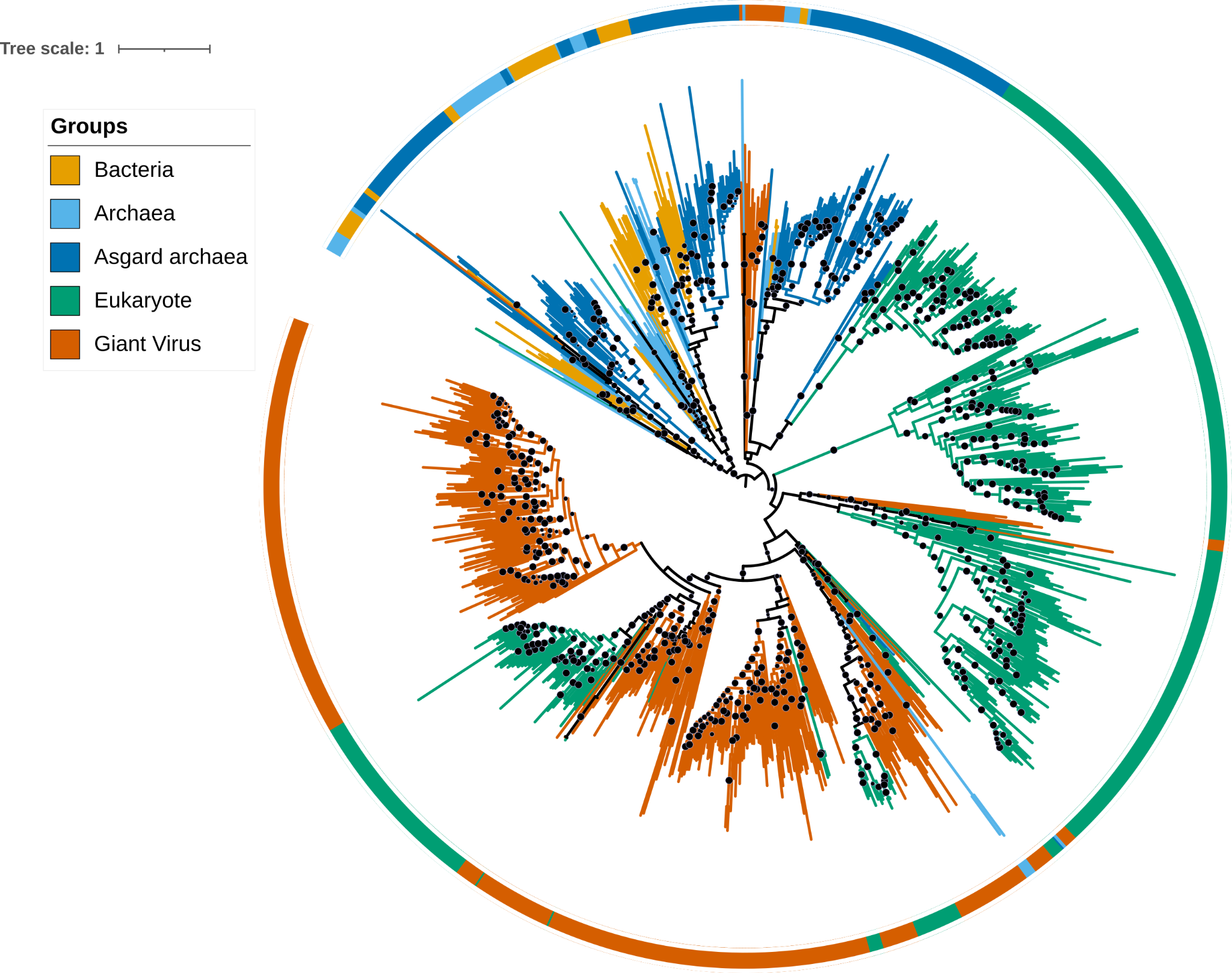

Figure 3: Polymerase family B phylogeny reconstructed using VT+F+R10 model (generated by -MFP parameter) with bootstrap support value represented by black dots ( bootstrap > 80% is shown).

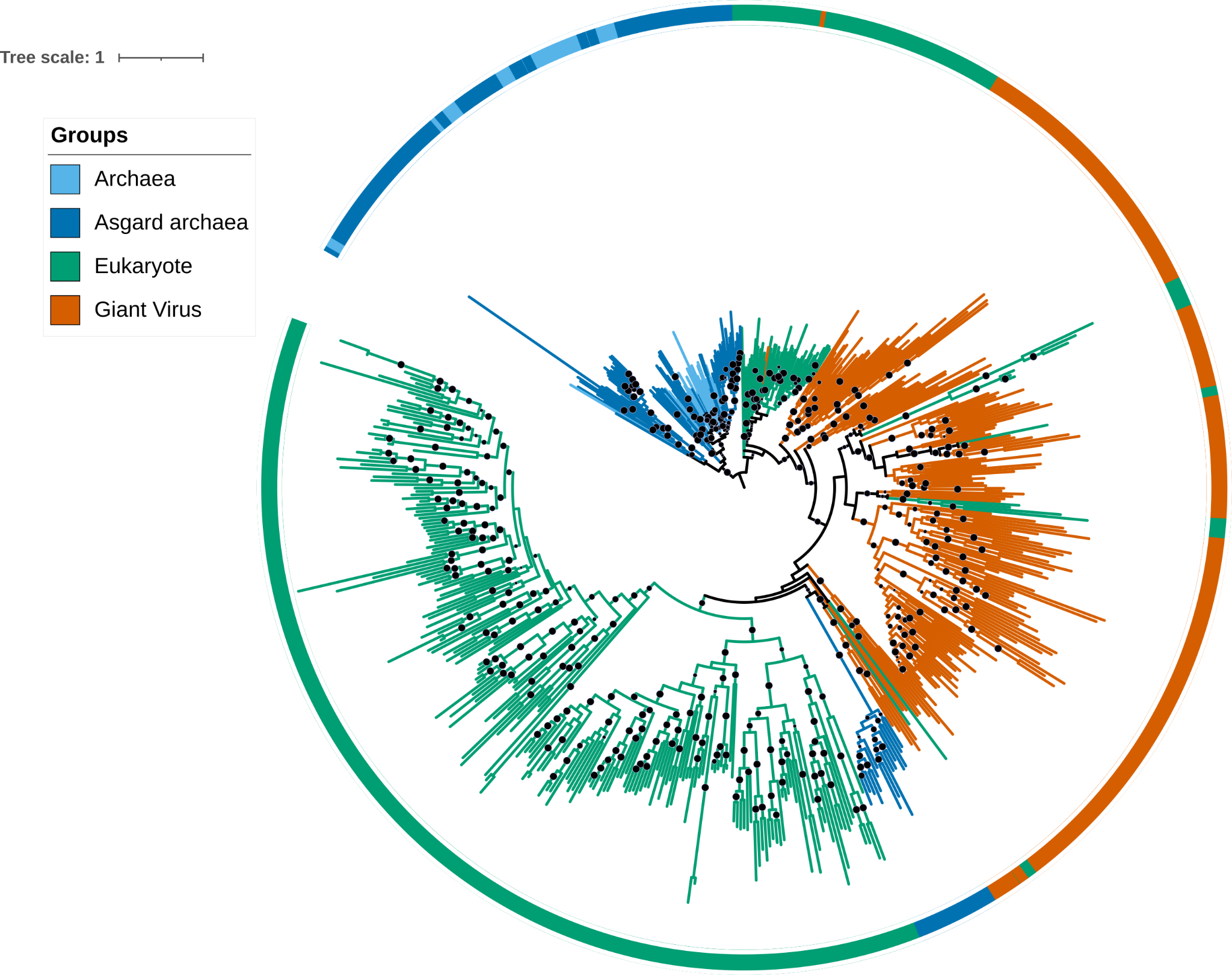

Figure 4: XPG endonuclease phylogeny reconstructed using VT+F+R10 model (generated by -MFP parameter) with bootstrap support value represented by black dots ( bootstrap > 80% is shown).

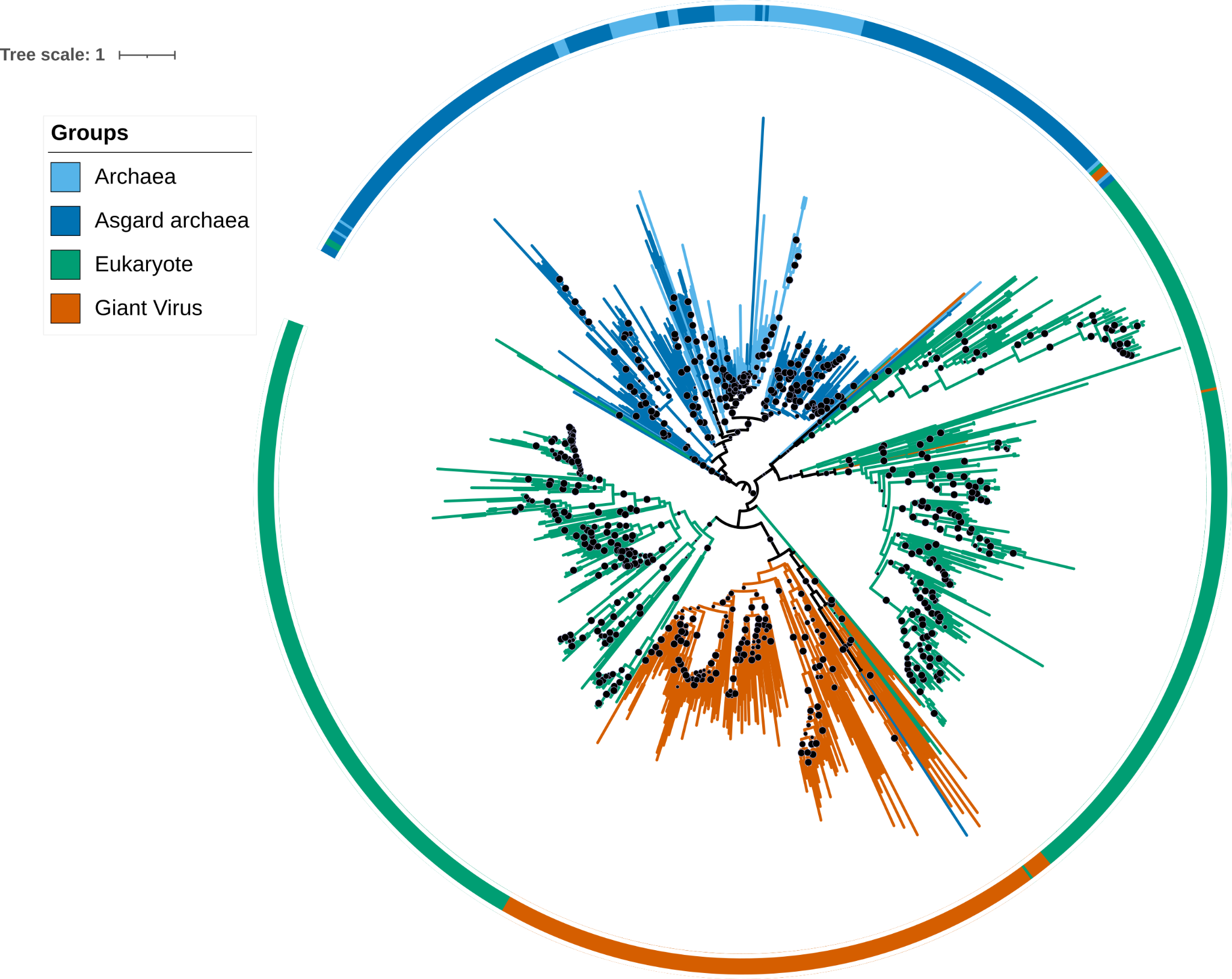

Figure 5: Transcription factor TFIIB phylogeny reconstructed using LG+R8 model (generated by -MFP parameter) with bootstrap support value represented by black dots ( bootstrap > 80% is shown).

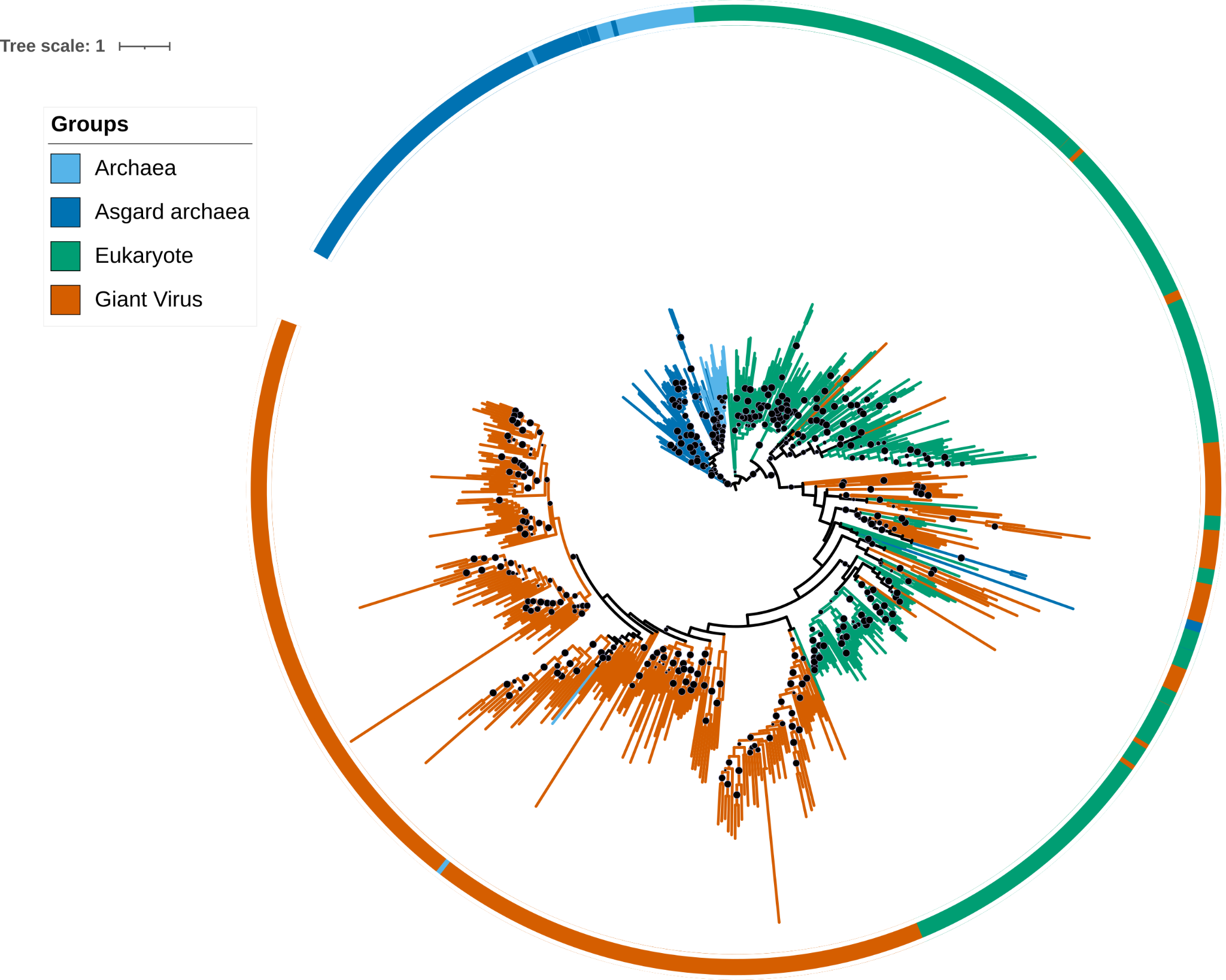

Figure 6: Transcription factor TFIIS phylogeny reconstructed using VT+F+R8 model (generated by -MFP parameter) with bootstrap support value represented by black dots ( bootstrap > 80% is shown).

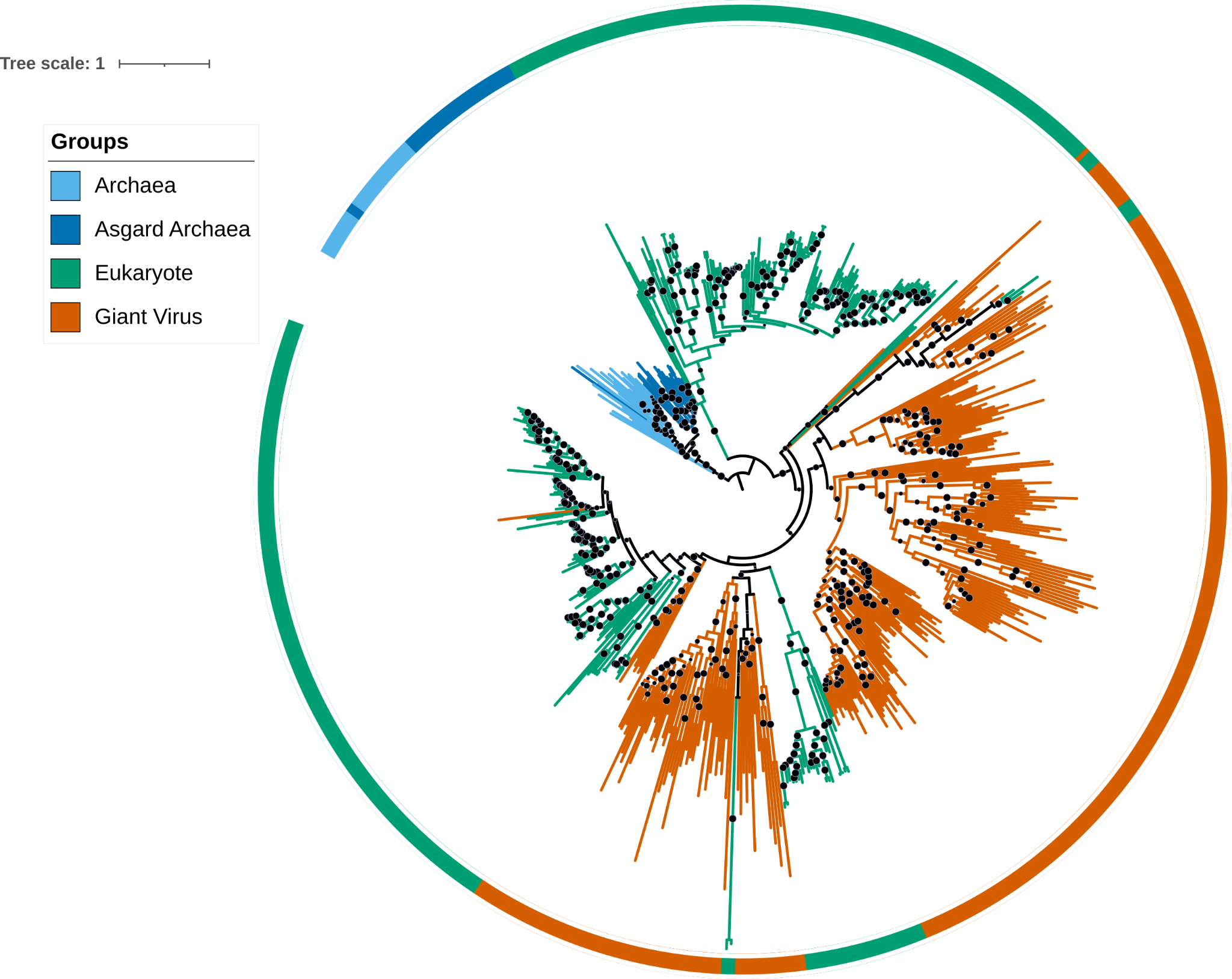

Figure 7: Phylogeny for RNA pol Rpb2, domain 4 reconstructed using LG+F+R10 model (generated by -MFP parameter) with bootstrap support value represented by black dots ( bootstrap > 80% is shown).

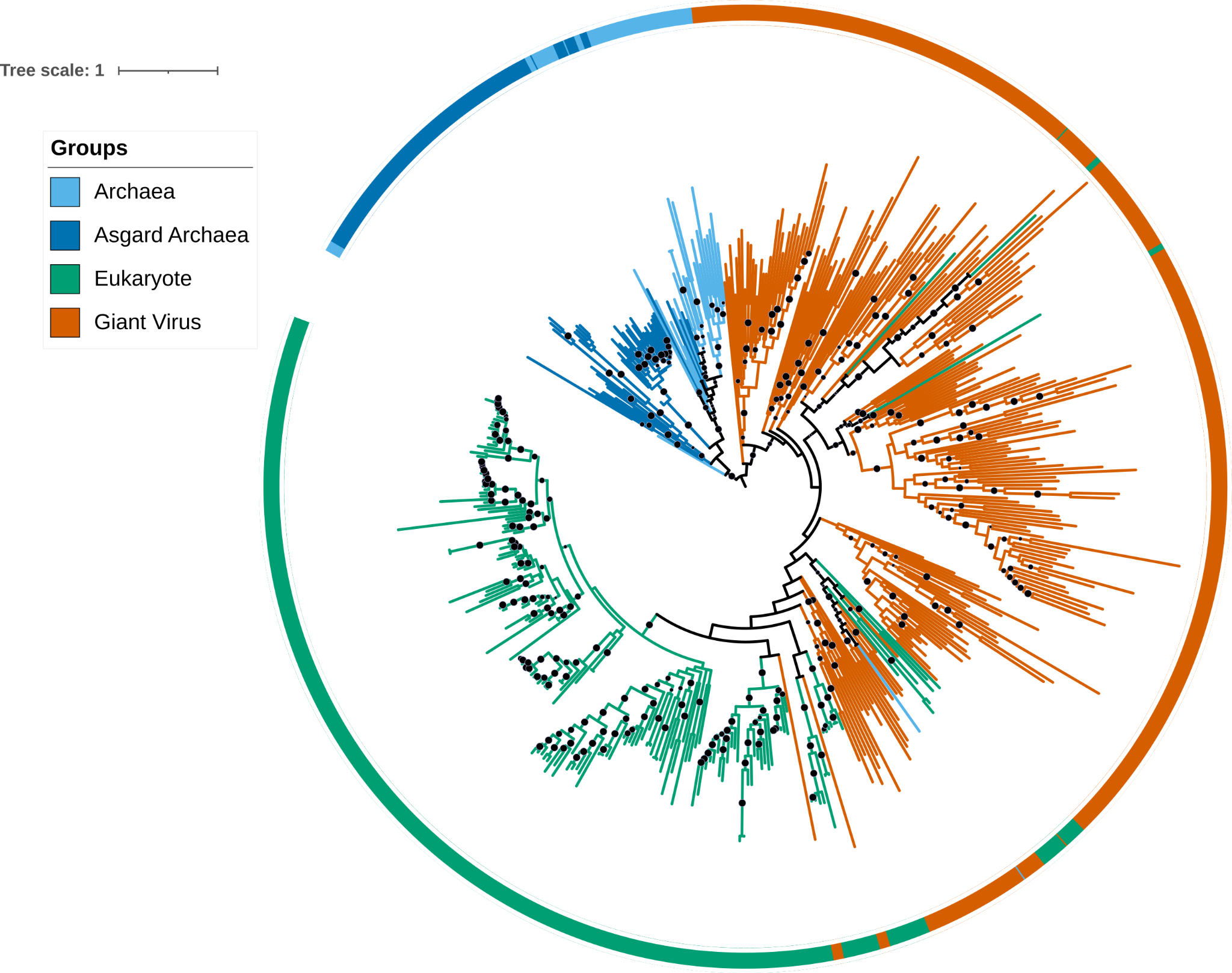

Figure 8: Phylogeny for RNA pol Rpb5, C terminal reconstructed using VT+F+R7 model (generated by -MFP parameter) with bootstrap support value represented by black dots ( bootstrap > 80% is shown).

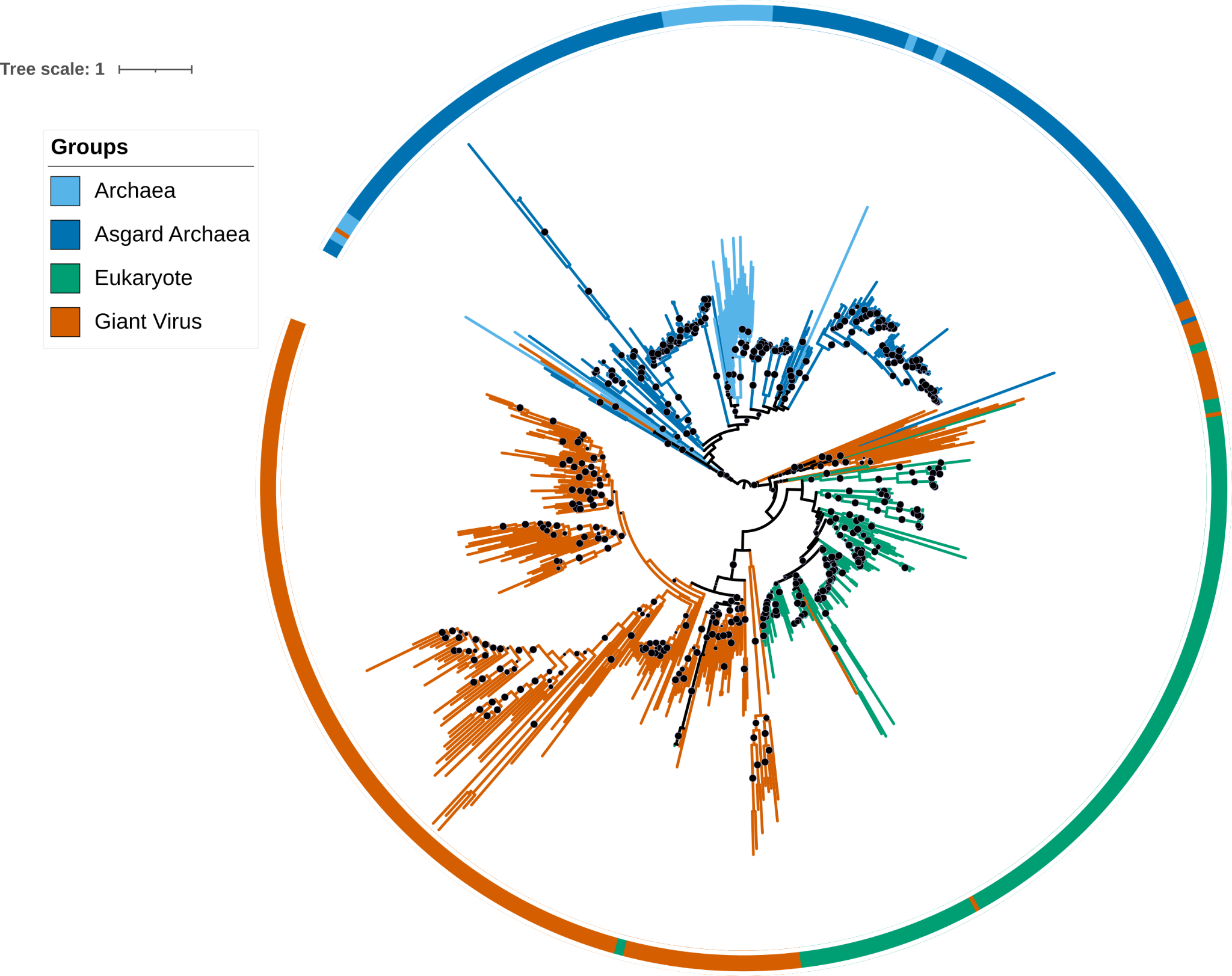

Figure 9: Phylogeny for DNA sliding clamp reconstructed using LG+F+R8 model (generated by -MFP parameter) with bootstrap support value represented by black dots ( bootstrap > 80% is shown).
