## Supplementary material for "Resolving ancient gene transfers clarifies the early co-evolution of eukaryotes and giant viruses": S3 Text

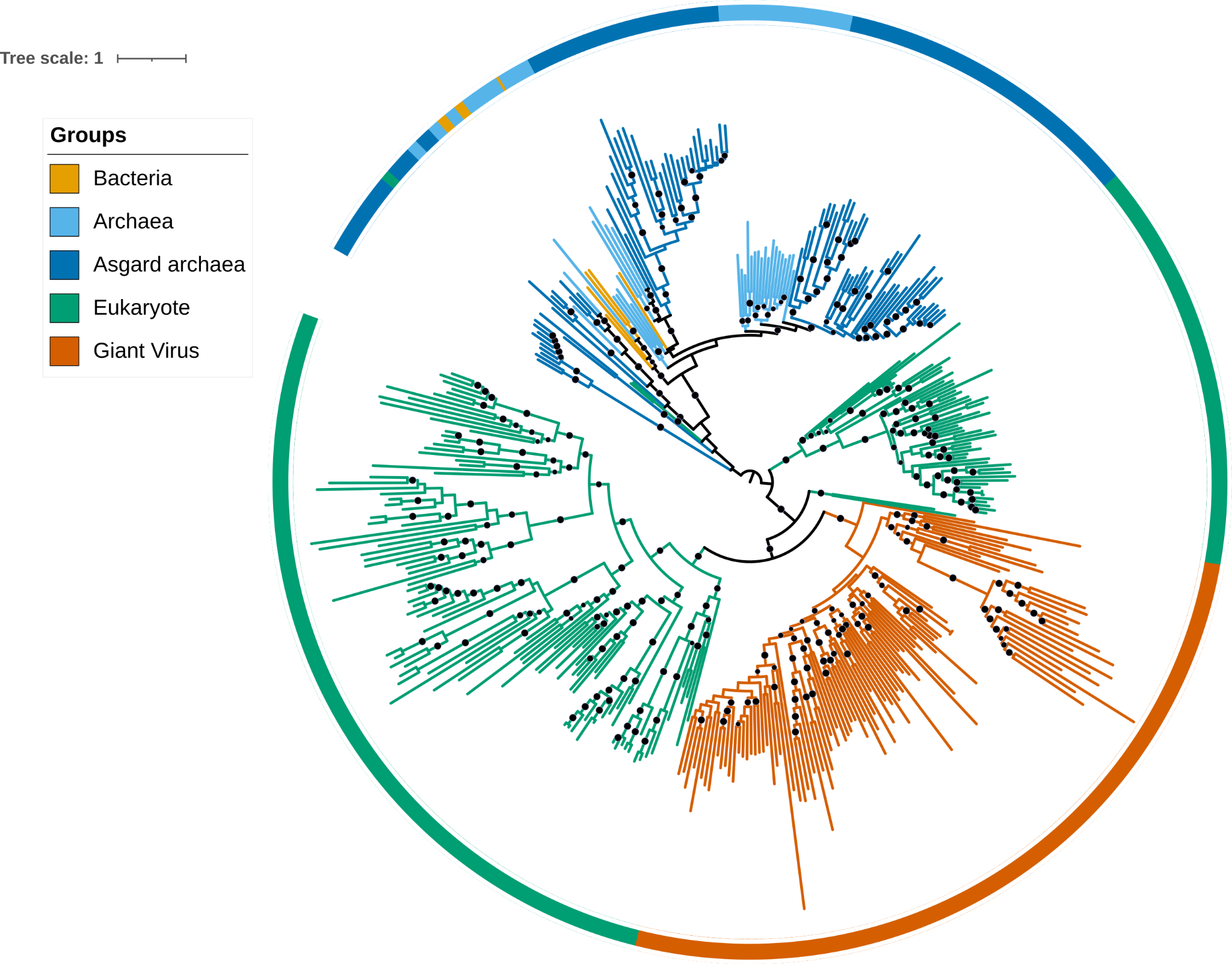

Figure 1: ERCC4 domain phylogeny reconstructed using a complex model (C60 + F +G) with bootstrap support value represented by black dots (bootstrap > 80% is shown).

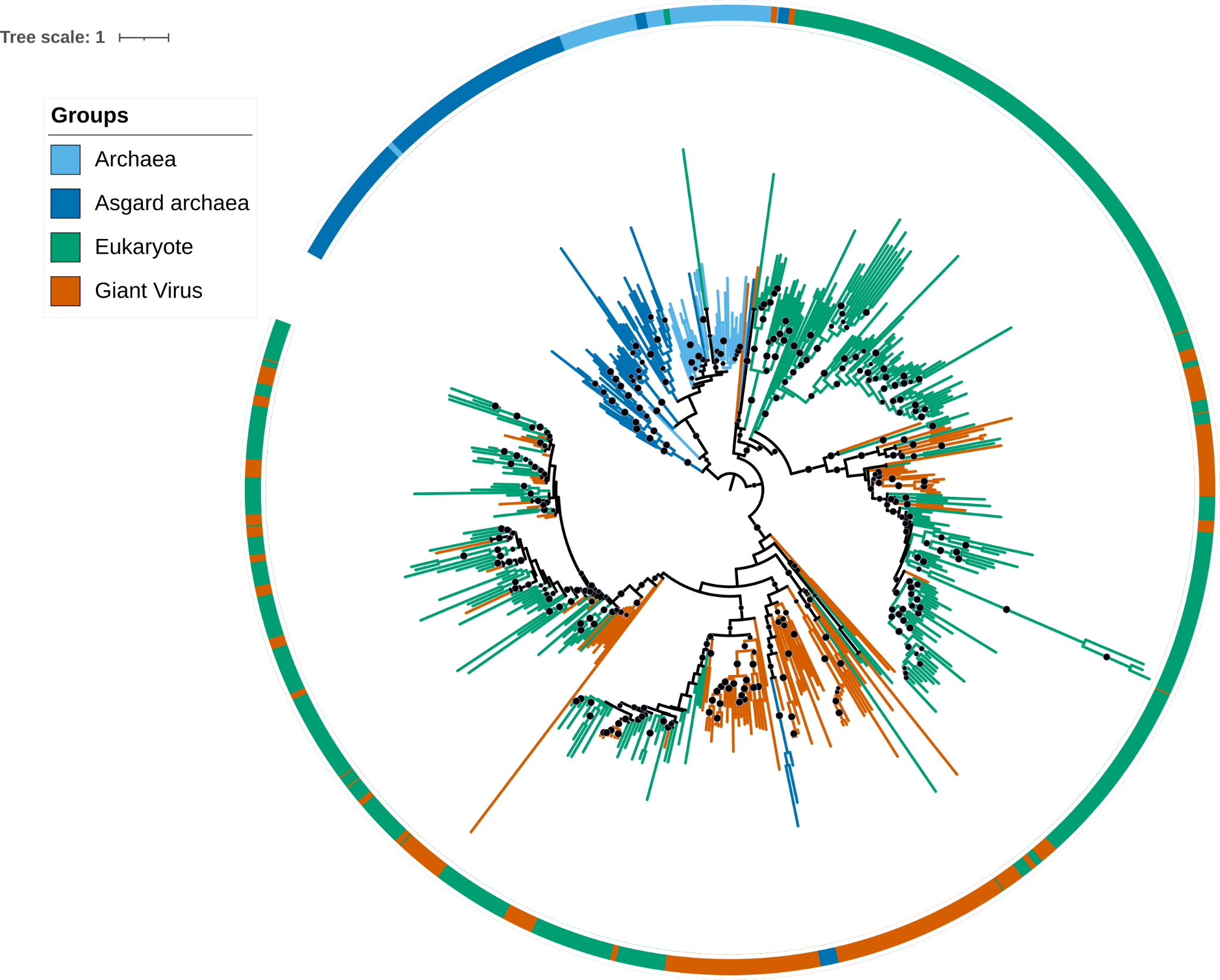

Figure 2: Core Histone phylogeny reconstructed using a complex model (C60 + F +G) with bootstrap support value represented by black dots (bootstrap > 80% is shown).

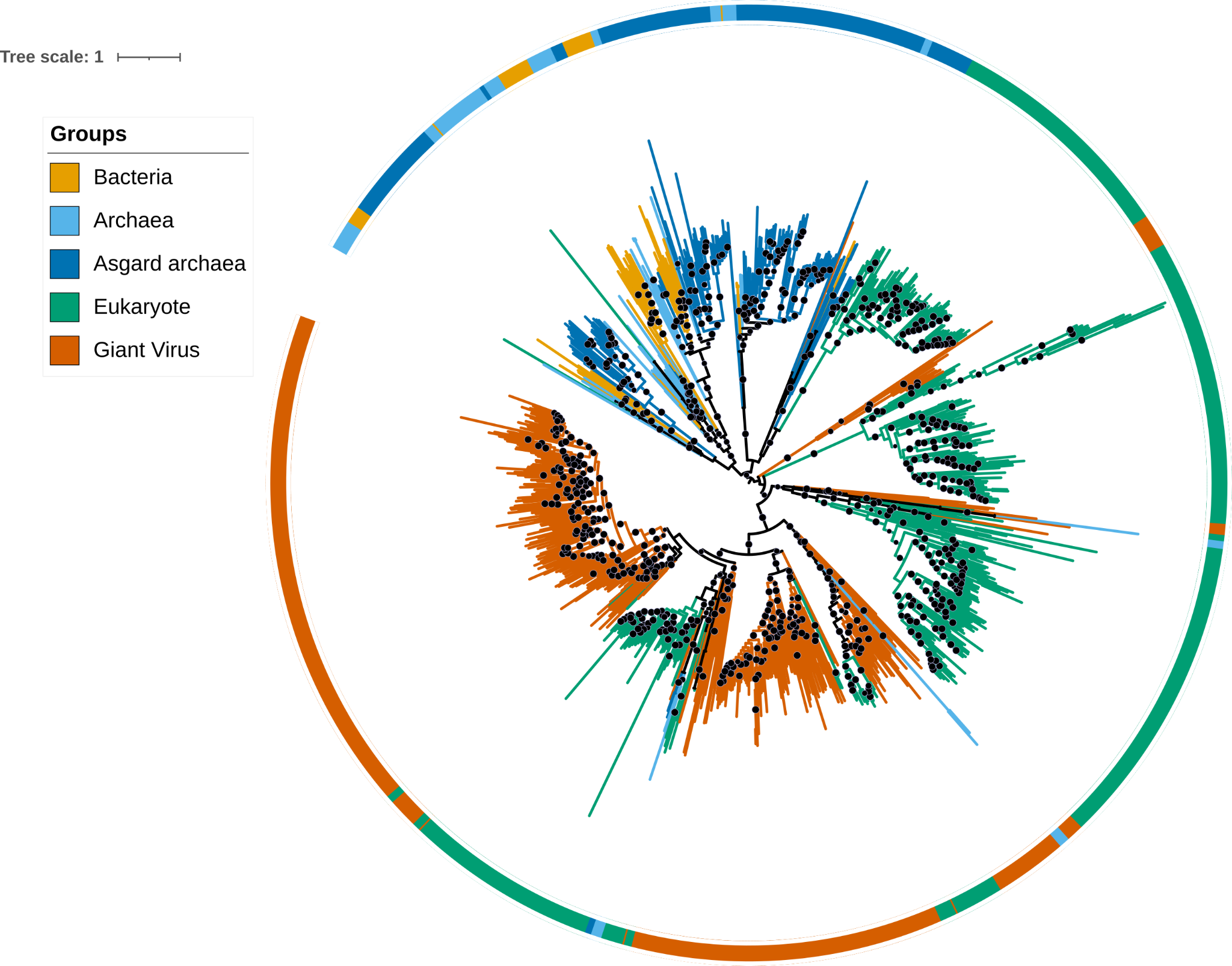

Figure 3: Polymerase family B phylogeny reconstructed using a complex model (C60 + F +G) with bootstrap support value represented by black dots (bootstrap > 80% is shown).

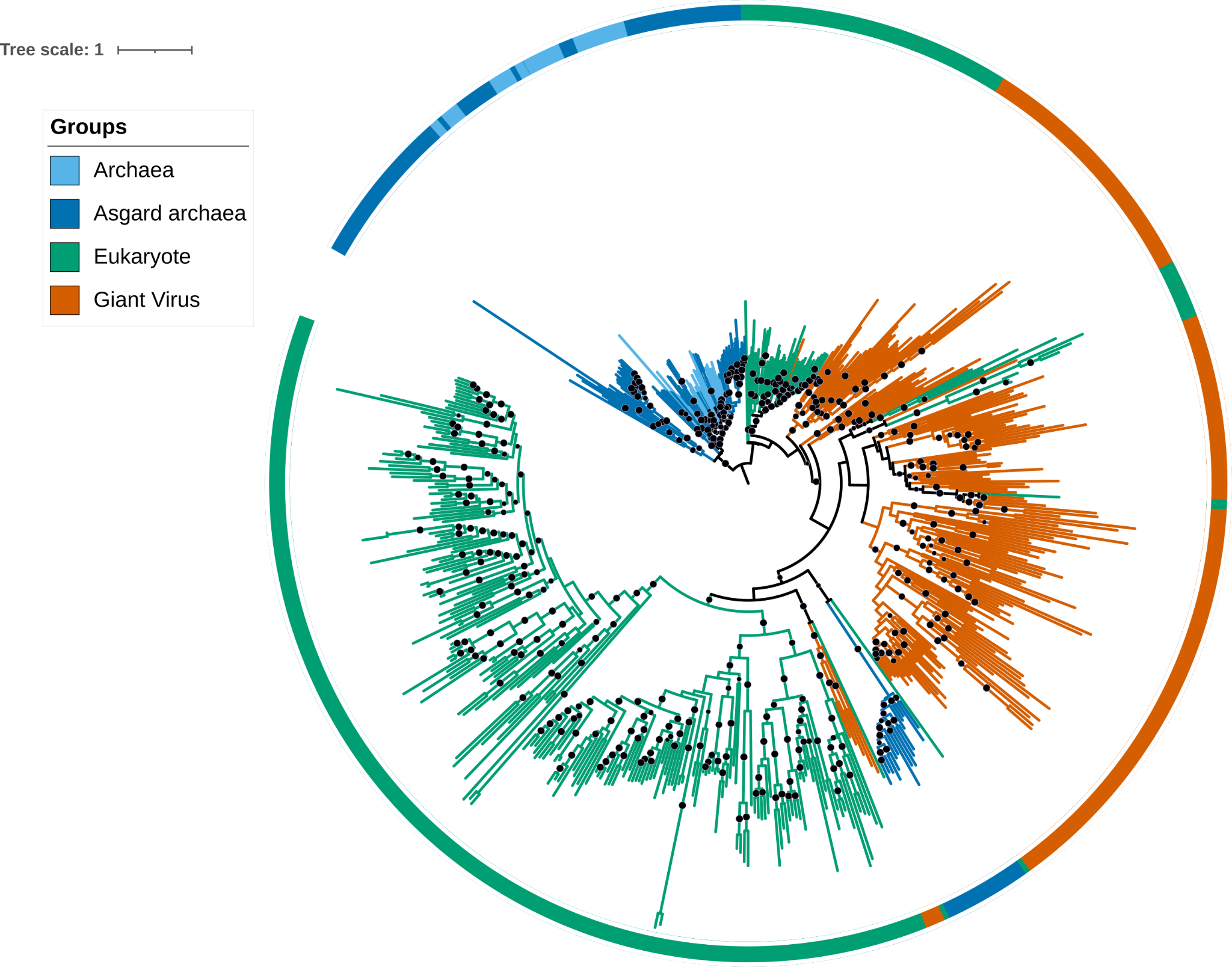

Figure 4: XPG endonuclease phylogeny reconstructed using a complex model (C60 + F +G) with bootstrap support value represented by black dots (bootstrap > 80% is shown).

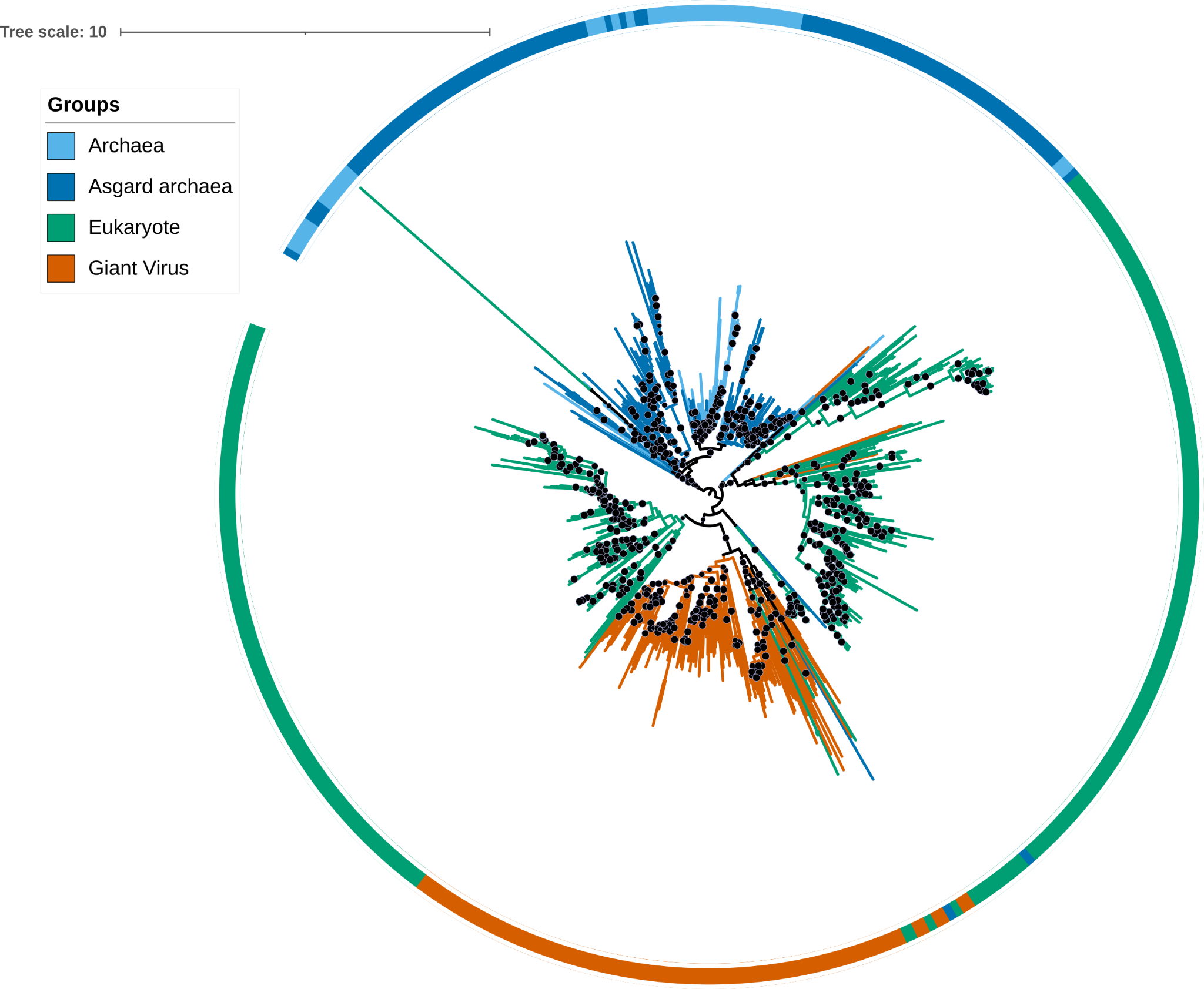

Figure 5: Transcription factor TFIIB phylogeny reconstructed using a complex model (C60 + F +G) with bootstrap support value represented by black dots (bootstrap > 80% is shown).

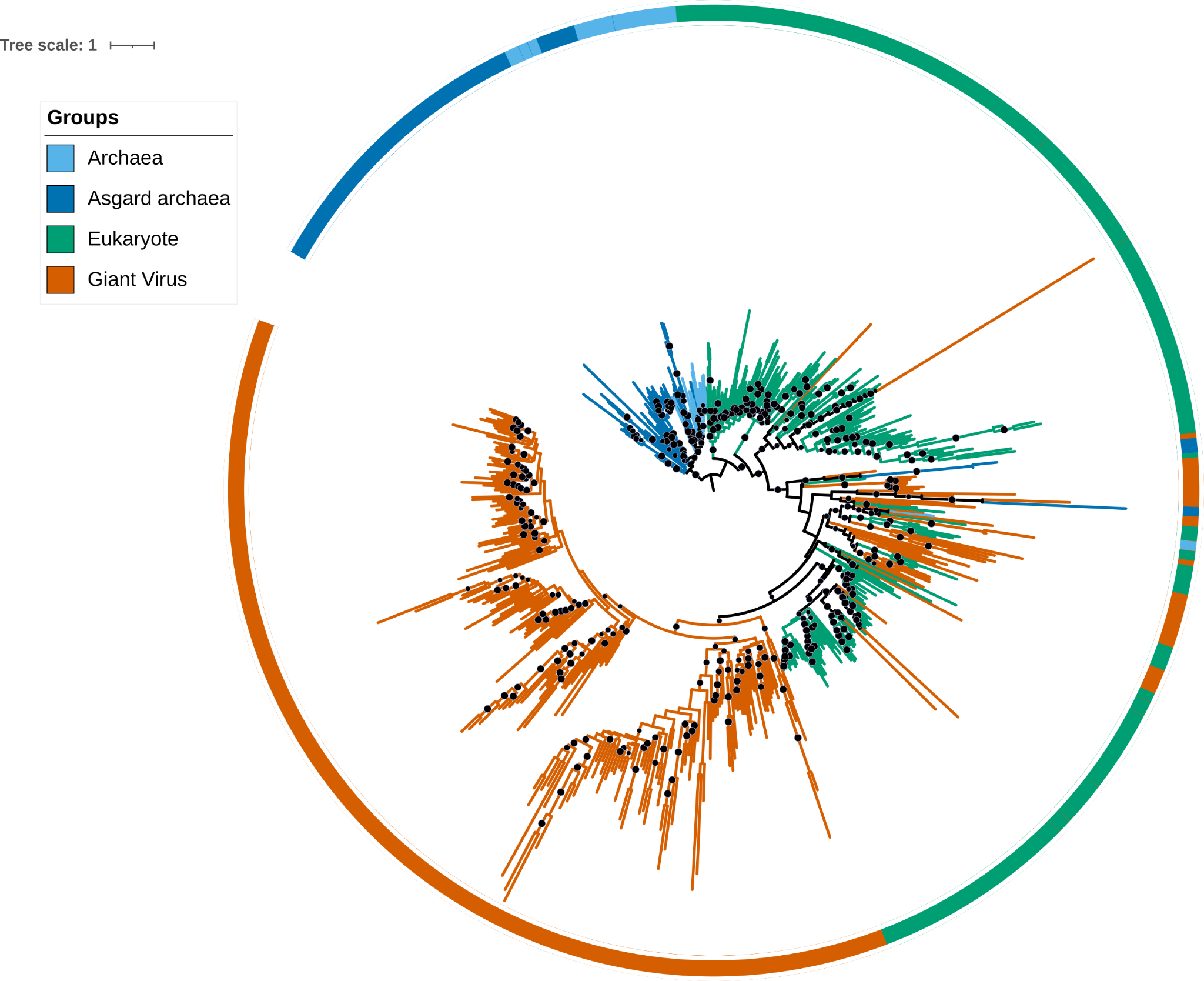

Figure 6: Transcription factor TFIIS phylogeny reconstructed using a complex model (C60 + F +G) with bootstrap support value represented by black dots (bootstrap > 80% is shown).

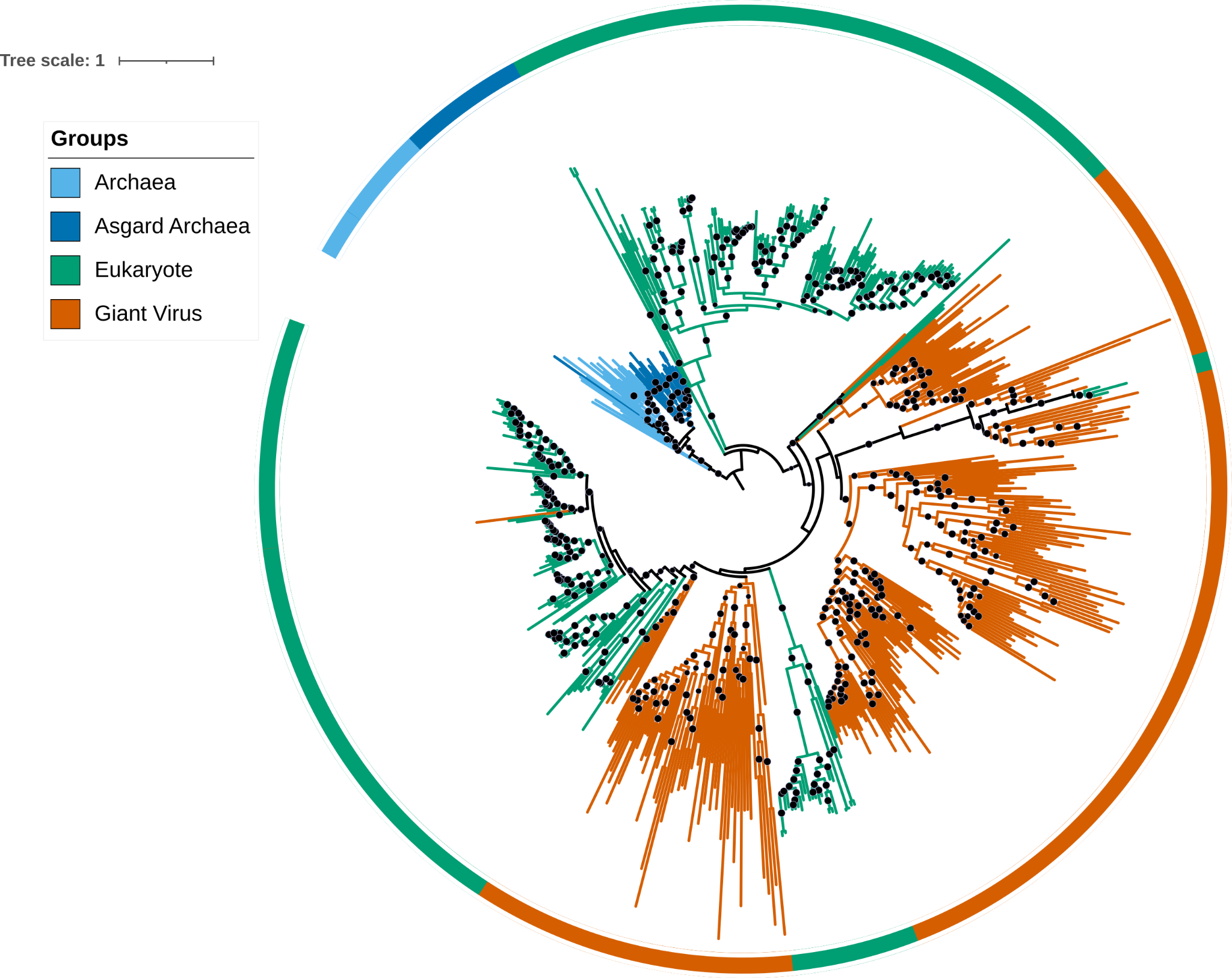

Figure 7: Phylogeny for RNA pol Rpb2, domain 4 reconstructed using a complex model (C60 + F +G) with bootstrap support value represented by black dots (bootstrap > 80% is shown).

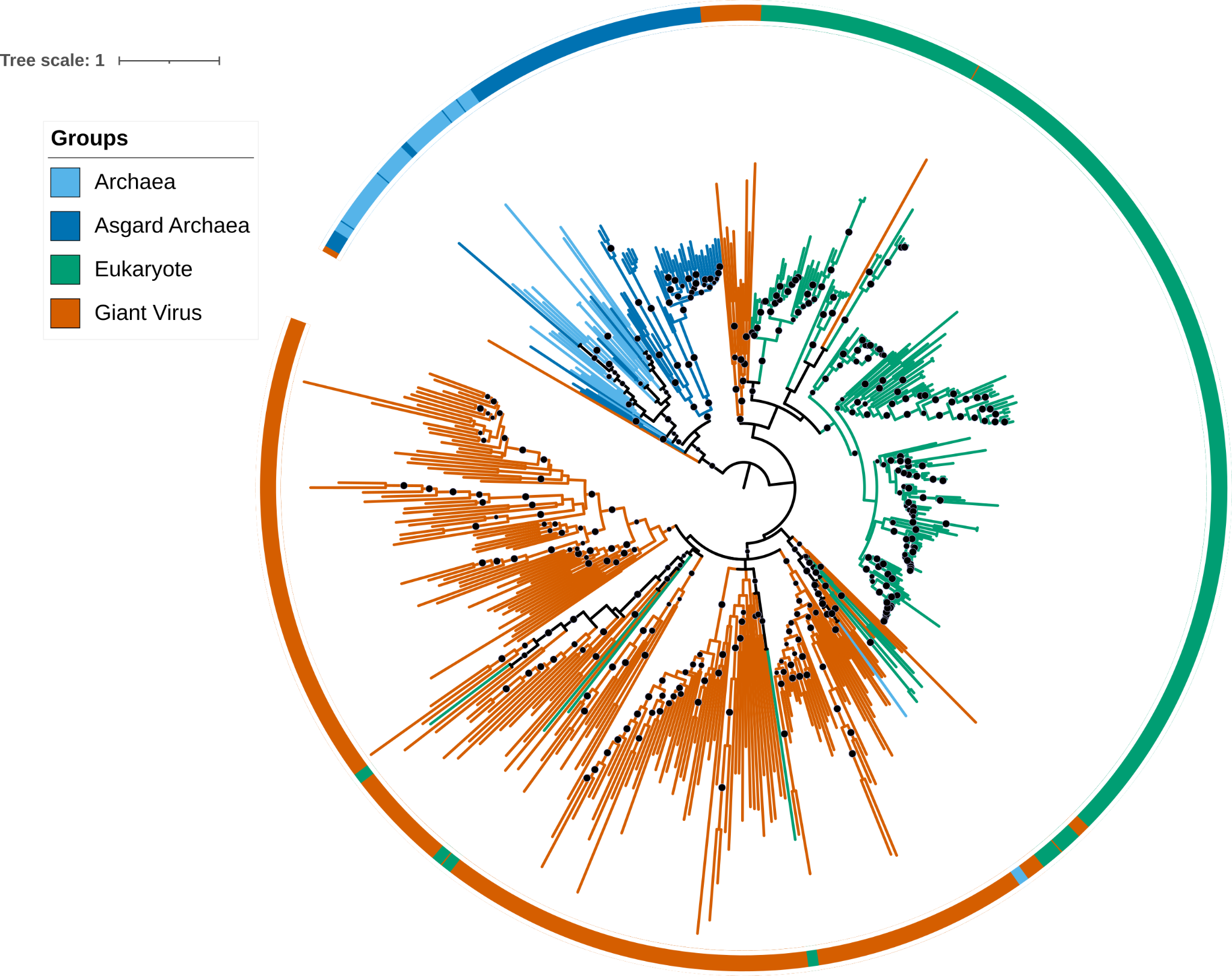

Figure 8: Phylogeny for RNA pol Rpb5, C terminal reconstructed using a complex model (C60 + F +G) with bootstrap support value represented by black dots (bootstrap > 80% is shown).

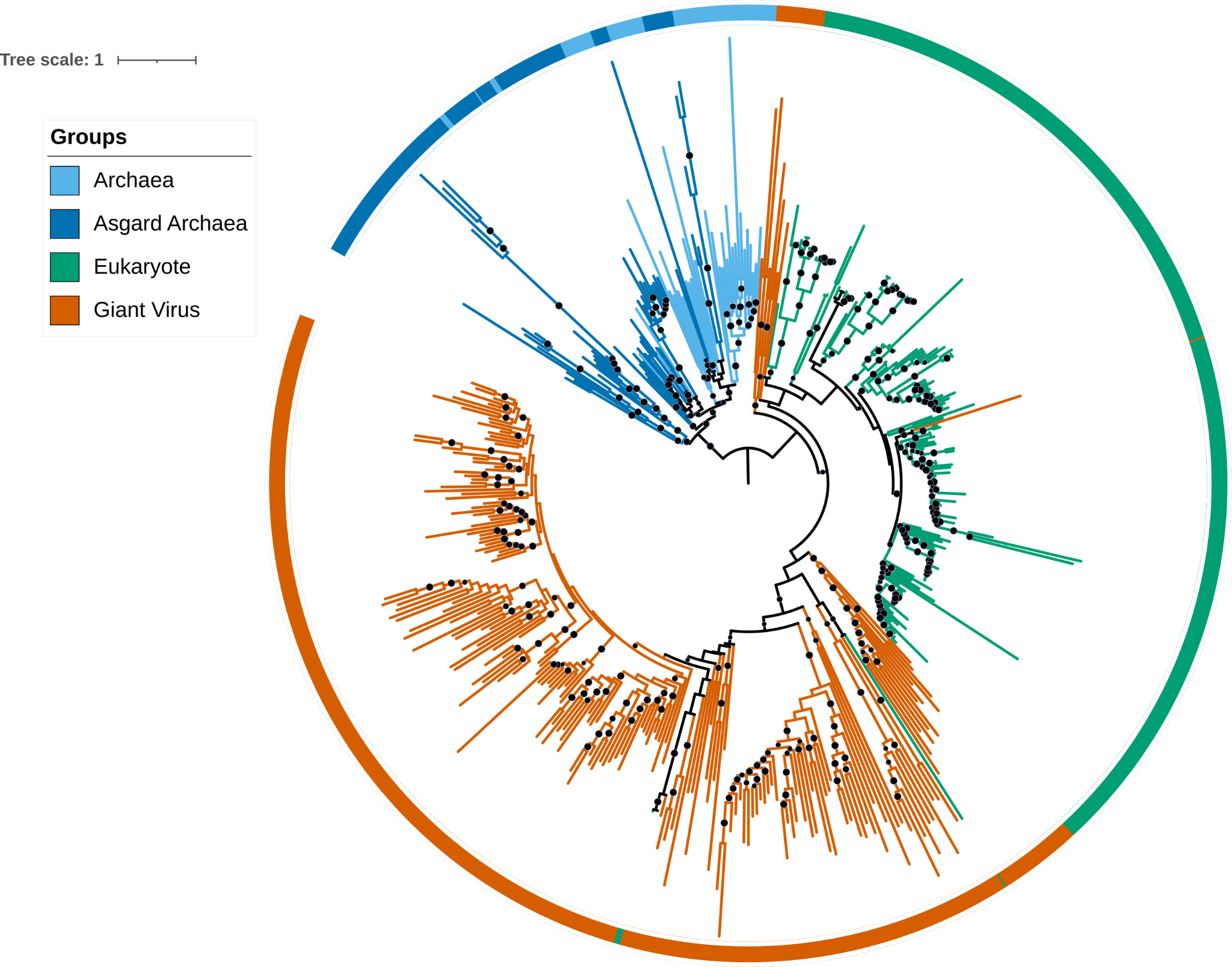

Figure 9: Phylogeny for PCNA reconstructed using a complex model (C60 + F +G) with bootstrap support value represented by black dots (bootstrap > 80% is shown).
